# Both GEF domains of the autism and epilepsy-associated Trio protein are required for proper tangential migration of GABAergic interneurons

**DOI:** 10.1101/2022.12.31.522400

**Authors:** Lara Eid, Ludmilla Lokmane, Praveen K. Raju, Samuel Boris Tene Tadoum, Xiao Jiang, Karolanne Toulouse, Alexis Lupien-Meilleur, François Charron-Ligez, Asmaa Toumi, Stéphanie Backer, Mathieu Lachance, Marisol Lavertu-Jolin, Marie Montseny, Jean-Claude Lacaille, Evelyne Bloch-Gallego, Elsa Rossignol

## Abstract

Recessive mutations in the *TRIO* gene are associated with intellectual deficiency (ID), autism spectrum disorder (ASD) and developmental epileptic encephalopathies (DEE). TRIO is a dual guanine nucleotide exchange factor (GEF) that activates Rac1, Cdc42 and RhoA. Trio has been extensively studied in excitatory neurons, and has recently been found to regulate the switch from tangential to radial migration in GABAergic interneurons (INs), through GEFD1-Rac1-dependent SDF1α/CXCR4 signalling. Given the central role of Rho-GTPases during neuronal migration and the implication of IN pathologies in ASD and DEE, we investigated the relative roles of both Trio’s GEF domains in regulating the dynamics of INs tangential migration. In *Trio^-/-^* mice, we observed reduced numbers of tangentially migrating INs, with intact progenitor proliferation. Further, we noted increased growth cone collapse in developing INs, suggesting altered cytoskeleton dynamics. To bypass the embryonic mortality of *Trio^-/-^* mice, we generated *Dlx5/6^Cre^;Trio^c/c^* conditional mutant mice, which develop spontaneous seizures and behavioral deficits reminiscent of ASD and ID. These phenotypes are associated with reduced cortical IN density and functional cortical inhibition. Mechanistically, this reduction of cortical IN numbers reflects a premature switch to radial migration, with an aberrant early entry in the cortical plate, as well as major deficits in cytoskeletal dynamics, including enhanced leading neurite branching and slower nucleokinesis reflecting reduced actin filament condensation and turnover. Further, we show that both Trio GEFD1 and GEFD2 domains are required for proper IN migration, with a dominant role of the RhoA-activating GEFD2 domain. Altogether, our data show a critical role of the DEE/ASD-associated *Trio* gene in the establishment of cortical inhibition and the requirement of both GEF domains in regulating IN migration dynamics.

## Introduction

Recessive and *de novo* mutations in the *TRIO* gene, encoding the triple functional domain protein, are associated with a spectrum of neurodevelopmental disorders (NDDs), including autism spectrum disorder (ASD), intellectual deficiency (ID) with microcephaly, and developmental epileptic encephalopathies (DEE) ^1–7^. Trio is a member of the large family of guanine nucleotide exchange factors (GEF), activators of small Rho-GTPases that promote the transition from an inactive GDP-bound to an active GTP-bound form of Rho-GTPases ^8^. Trio is composed of two distinct Rac-specific and Rho-specific guanine nucleotide exchange factor (GEF) domains as well as a serine/threonine kinase domain ^9^. In neurons, Trio’s GEFD1 activates Rac1, RhoG and Cdc42 ^10–14^ while GEFD2 activates RhoA ^14, 15^. Trio thus acts as a molecular switch that controls multiple small Rho-GTPases in response to a variety of environmental cues relevant to the migration and morphological maturation of neurons.

Recent functional validation of *TRIO* mutations in either heterologous cellular systems, cultured neurons or mutant mouse brains suggest that most ASD/ID-associated variants, particularly those in the GEFD1 domain, reduce TRIO-GEFD1-dependant Rac1 activation ^2–4, 6, 7, 16^, although a few *de novo* sporadic variants result in an aberrant hyperactivation of Rac1 ^4^. However, the network mechanisms by which *TRIO* mutations result in NDDs are unclear.

In mammals, Trio expression is ubiquitous and vital as the constitutive loss of *Trio* is embryonically lethal ^17^. However, studies in knockout mice indicate major roles of Trio in regulating the cellular organization of the hippocampus and olfactory bulb ^17^, the formation of neuronal clusters in the hindbrain ^18^, the migration and morphology of cerebellar granule cells ^13^, as well as the guidance of cortico-spinal axons ^10^ and of thalamocortical axons through its regulation of corridor cell migration ^15^. Furthermore, Trio controls the morphological maturation and function of excitatory neurons ^1, 4, 7, 19^ and was recently found to participate in the migration of GABAergic interneurons (INs) ^16^. At the molecular level, Trio GEFD1 regulates both cytoskeletal dynamics ^13, 15^ and membrane trafficking of the Trio protein ^20^, regulating axonal guidance and neurite outgrowth through netrin-1 induced activation of the Rac1/RhoG and Cdc42 pathways ^10, 12, 14^. Furthermore, the GEFD2 domain controls axonal pathfinding through Slit2-induced activation of the RhoA pathway ^15^.

Given the embryonic lethality of *Trio^-/-^* mice ^17^, conditional mouse models have been used to study the impact of *Trio* deletions on specific neuronal populations. The targeted deletion of *Trio* in neural progenitor cells, using the *Nestin-Cre* driver line ^21^, results in early perinatal mortality and ataxia in survivors ^13^, mostly through a GEFD1-dependent cerebellar granule cell migration deficit ^14^. The targeted deletion in cortical pyramidal cells, using the *Emx1-Cre* driver line ^22, 23^, results in a milder phenotype, with intact survival but abnormal hippocampal organization leading to spatial learning deficits ^24^, as well as impaired morphogenesis and migration of late-born cortical pyramidal cells ^25^. Interestingly, the targeted deletion of *Trio* in INs was recently shown to induce an ASD-like phenotype ^16^, although the molecular, cellular and network mechanisms underlying this neurodevelopmental phenotype remain unclear.

Here, we show that the constitutive deletion of *Trio* impairs the tangential migration of INs, without affecting the proliferation of IN progenitors. Using a conditional genetic strategy, we show that the specific prenatal deletion of *Trio* in GABAergic INs impairs cytoskeletal remodeling in tangentially migrating INs, with major contributions of the GEFD2-RhoA pathway, resulting in delayed tangential migration, reduced cortical inhibition, epilepsy and cognitive-behavioral deficits. Together, our data support a critical role for *Trio* and the regulation of Rho-GTPases during IN development in the pathogenesis of NDD, including ASD and DEE.

## Results

### Trio is critical for the migration of GABAergic INs

To investigate whether *Trio* plays a role in the early development and migration of GABAergic INs, we first quantified dorsally-migrating *Lim homeobox 6* (*Lhx6*)-expressing INs in *Trio^-/-^* knockout embryos using *in situ* hybridization to selectively label the sub-populations derived from the medial ganglionic eminence (MGE). We found a reduction of *Lhx6*-positive (+) INs entering the dorsal pallium at e14.5 in *Trio^-/-^* mutants compared to wild-type (WT) embryos, leading to a disrupted laminar distribution of *Lhx6+* GABAergic INs at e18.5 in *Trio^-/-^* brains, compared to WT littermates (Supplementary Figure 1 A-F).

Such a reduction in migrating INs could reflect a primary disruption of neuronal proliferation. To investigate this possibility, we pulse-labeled e12.5 embryos with 5-ethynyl-2’-deoxyuridine (EdU) and examined EdU incorporation in neural progenitor cells 2h after injection. We found similar numbers of dividing MGE progenitors in *Trio^-/-^* mutants and WT littermates, suggesting that the deletion of *Trio* does not impair the proliferation of MGE-derived IN progenitors (Supplementary Figure 2).

To better characterize the cell-autonomous mechanisms by which IN migration is impaired in *Trio^-/-^* knockout embryos, we generated MGE explants from e14.5 mouse embryos that were cultured in collagen for 48h. We analyzed both the cell body migration and leading processes extensions. While both WT and *Trio^-/-^* MGE-cells had the capacity to migrate out of the MGE explants, we found that migrating *Trio^-/-^* MGE-cells traveled a shorter distance compared to WT MGE-cells (Supplementary Figure 1 G-K). Furthermore, *Trio^-/-^* MGE-cells in dissociated cultures exhibited an increased rate of growth cone collapse, while growth cones with a complex morphology occupied a larger surface than in WT MGE-cells (Supplementary Figure 1 L-Q). These results suggest that the loss of *Trio* impairs the migration of GABAergic INs in a cell-autonomous fashion.

### Trio deletion in GABAergic INs induces epilepsy as well as cognitive and behavioral deficits

To further study the requirement of *Trio* in INs and its relevance to the overall clinical phenotype, while bypassing the embryonic lethality of *Trio^-/-^* mice, we generated conditional knock-out mice (*Trio^cKO^*) carrying a targeted deletion of *Trio* in GABAergic INs. Specifically, we bred *Trio^lox/lox^* mice, in which exons 22-25 are flanked by two *loxp* sites ^13^, with *Dlx5/6^Cre^;RCE^EGFP^* mice, to selectively target telencephalic GABAergic INs^26, 27^, while labeling Cre-expressing cells in green with the *RCE^EGFP^* reporter allele ^28^.

*Trio^cKO^* animals are viable, although mutant mice appear smaller than their *Trio^WT^* littermates. Notably, *Trio^cKO^* mice develop spontaneous seizures as early as P14, and ≈50% of mutant mice die before P30. To better characterize the seizure phenotype, we recorded video-EEG for 72h in surviving *Trio^cKO^* mice at P30-P36 (Figure 1 A-B). These recordings revealed abundant interictal epileptic activity in all mutants, as well as spontaneous seizures in 3 out of 4 mutants recorded (Figure 1 B), with a mean seizure severity score of 3.4 ± 0.3 on the modified Racine scale (Figure 1 C), a mean seizure frequency of 2.0 ± 0.9 seizures per 72h (Figure 1 D) and a mean seizure duration of 22.5 ± 2.3 seconds (Figure 1 E).

**Figure 1.**
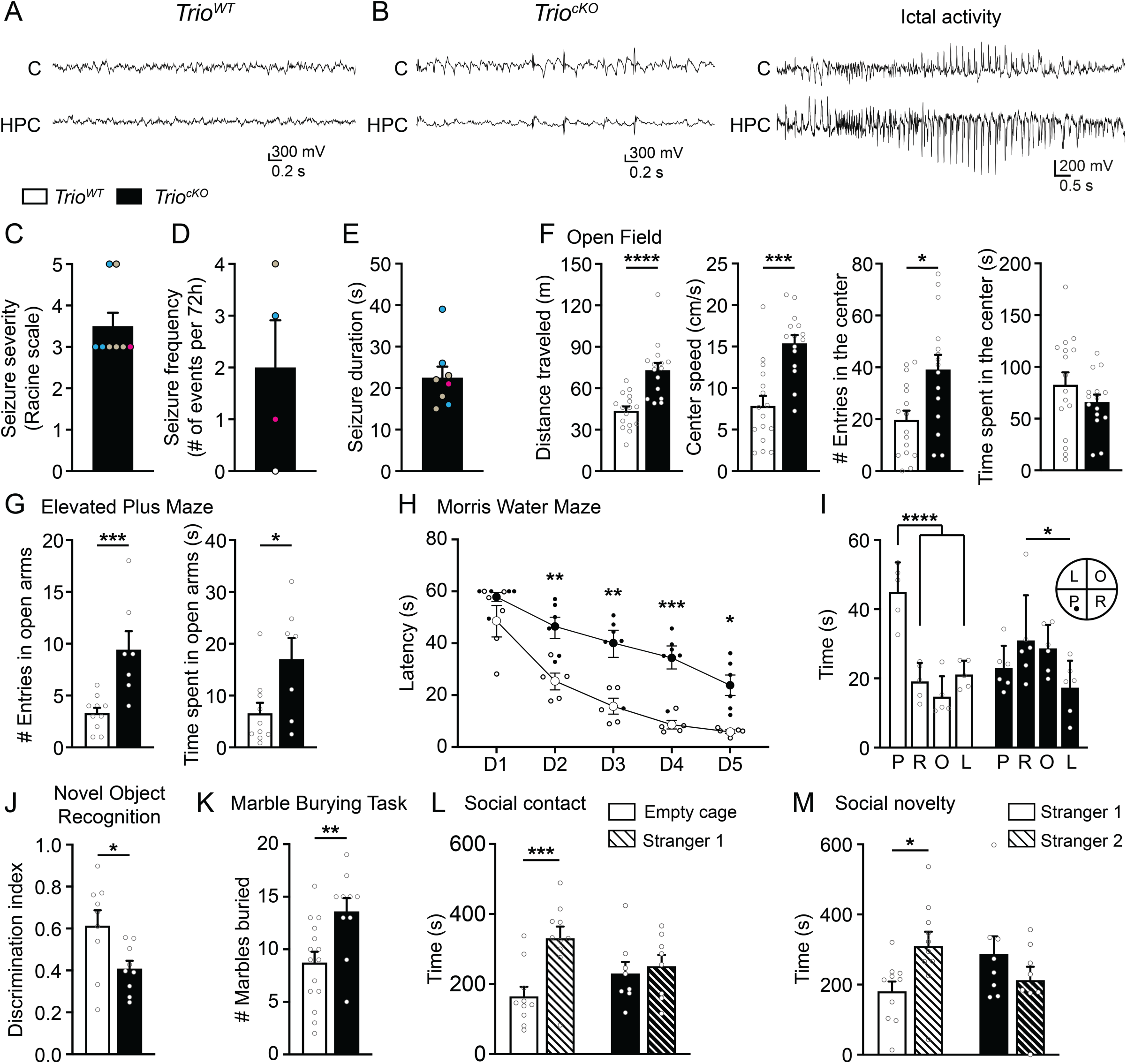
Conditional deletion of *Trio* in GABAergic INs leads to seizures and cognitive-behavioral deficits. (**A-E**) Field potentials and seizure activity assessed with video-EEG recordings in P30-36 littermates. (**A**) Representative EEG traces for *Trio^WT^* and (**B**) *Trio^cKO^* mice, showing abundant inter-ictal activity (left traces) and a generalized tonic-clonic seizure (right traces) in mutants. Recordings were performed using bilateral implanted cortical (C) and hippocampal (HPC) electrodes. For illustration purposes, recordings from single cortical (C) and hippocampal (HPC) fields are illustrated. (**C-E**) Histograms displaying the seizure severity (**C**), frequency (**D**) and duration (**E**) for each animal (color-coded for individual mice) (n = 4 *Trio^WT^* and 4 *Trio^cKO^*). (**F-M**) Behavioral deficits in *Trio^cKO^* mice. (**F**) Histograms showing the distance traveled, the mean speed, the number of entries and the time spent in the center zone during the open field test (n = 16 *Trio^WT^* and 15 *Trio^cKO^* mice. (**G**) Histograms showing the number of entries in the open arms (left) and the time spent in the open arms (right) in the elevated plus maze (n = 10 *Trio^WT^* and 7 *Trio^cKO^* mice). (**H**) Histogram showing the escape latency in the Morris water maze during the acquisition phase (Day (D) 1 to D5). (**I**) Histogram showing the time spent in each quadrant of the Morris water maze during the probe test (P: platform quadrant, R: adjacent right quadrant, O: opposite quadrant, L: adjacent left quadrant; N = 5 *Trio^WT^* and 6 *Trio^cKO^* mice). (**J**) Histogram showing the discrimination index for a novel object over a familial object during the novel object recognition task (N = 9 *Trio^WT^* and 9 *Trio^cKO^* mice). (**K**) Histogram showing the number of marbles buried during the Marble burying task (N = 15 *Trio^WT^* and 10 *Trio^cKO^* mice). (**L**) Histogram showing the time spent with an empty cage or with another mouse (stranger 1) during the socialization phase of the three-chamber maze task and (**M**) Histogram showing the time spent with a familiar mouse (stranger 1) and a novel mouse (stranger 2) in the novelty phase of the three-chamber maze task (N = 10 *Trio^WT^* and 8 *Trio^cKO^* mice). \**P* < 0.05, \*\**P* < 0.01, \*\*\**P* < 0.001 and \*\*\*\**P* < 0.0001, by Mann-Whitney test (**F, G, J, K**), two-way ANOVA followed by Bonferroni’s multiple comparisons test (**H, L, M**) or two-way ANOVA followed by Tukey’s multiple comparisons test (**I**).

To further characterize the clinical phenotype of *Trio^cKO^* mice, we next examined their behavior in various paradigms. In the Open field, *Trio^cKO^* mice displayed frank motor hyperactivity, with longer distances travelled and increased speed compared to *Trio^WT^* (Figure 1 F). Further, *Trio^cKO^* mice displayed reduced anxiety in the Open Field, with increased numbers of entries in the center zone of the maze (Figure 1 F). This reduced fear of open spaces was confirmed in the Elevated plus maze, as *Trio^cKO^* mice entered more frequently and spent more time in the open arms than their *Trio^WT^* littermates (Figure 1 G). Further, in the Morris water maze, *Trio^cKO^* mice displayed striking spatial learning deficits, with increased escape latency in days 2-5 compared to *Trio^WT^* (Figure 1 H). *Trio^cKO^* mice also displayed reduced spatial memory in the probe test: they spent as much time in all quadrants as in the target platform quadrant compared to *Trio^WT^* who spent more time in the target platform quadrant (Figure 1 I). In the novel object recognition task, *Trio^cKO^* mice failed to adequately discriminate between novel and familiar objects (Figure 1 J), suggesting a reduction in novelty seeking in *Trio^cKO^* mice compared to *Trio^WT^*. In addition, *Trio^cKO^* mice presented increased repetitive behaviors in the marble-burying task: *Trio^cKO^* animals buried nearly twice as many marbles as their *Trio^WT^* littermates (Figure 1 K). Finally, *Trio^cKO^* mice presented reduced socialization skills in the three-chamber maze, spending as much time with an empty cage as with a live animal, as opposed to WT mice who clearly prefer interacting with a live animal (stranger 1; Figure 1 L). *Trio^cKO^* mice also spent as much time with a novel mouse (stranger 2) as with a familiar mouse (stranger 1; Figure 1 M), indicating a deficit in social novelty seeking. Taken together, these results suggest that the deletion of *Trio* in GABAergic INs induces an ASD and DEE-like phenotype in mice, with hyperactivity, reduced anxiety (risk-taking behaviors), impaired novelty seeking, socialization deficits and increased repetitive behaviors, as well as spatial learning and memory deficits and spontaneous seizures.

### Trio deletion disrupts the final laminar distribution of cortical GABAergic INs resulting in reduced cortical inhibition

To assess whether the targeted loss of Trio impacts the final numbers of cortical INs in *Trio^cKO^* mice, we quantified fate-mapped cortical GABAergic INs in the somatosensory cortex (S1) of P21 *Trio^cKO^* and *Trio^WT^* mice (Figure 2 A, B). We found a 15% net reduction of GFP+ cells in S1, particularly in layer I (23%), layer II-III (28%) and layer VI (20%) of *Trio^cKO^* mice compared to *Trio^WT^* littermates (Figure 2 C, D). To assess which IN sub-populations were affected, we quantified the number of fate-mapped GFP+ INs that expressed either parvalbumin (PV) (Figure 2 E, F), somatostatin (SST) (Figure 2 I, J), calretinin (CR) (Figure 2 M, N), or vaso-intestinal peptide (VIP) (Figure Q, R). We found a 20% decrease in the density of PV-expressing INs in S1, particularly in layers V (22%) and VIP (45%) of *Trio^cKO^* mice, compared to *Trio^WT^* littermates (Figure 2 G, H). By contrast, SST-expressing INs numbers remained unchanged in all cortical layers (Figure 2 K, L). Further, we found a 28% decrease in the density of CR+ INs in S1 of *Trio^cKO^* mice, which was particularly evident in layer II-III (31%) and layer IV (33%), when compared to *Trio^WT^* littermates (Figure 2 O, P). We also found a 27% reduction in the density of VIP+ INs in layer II-III (Figure 1 S, T). Thus, the loss of Trio affects multiple populations of INs, with striking reductions in PV-expressing INs and in CR- (but not SST)-expressing INs, as well as in VIP-expressing INs.

**Figure 2.**
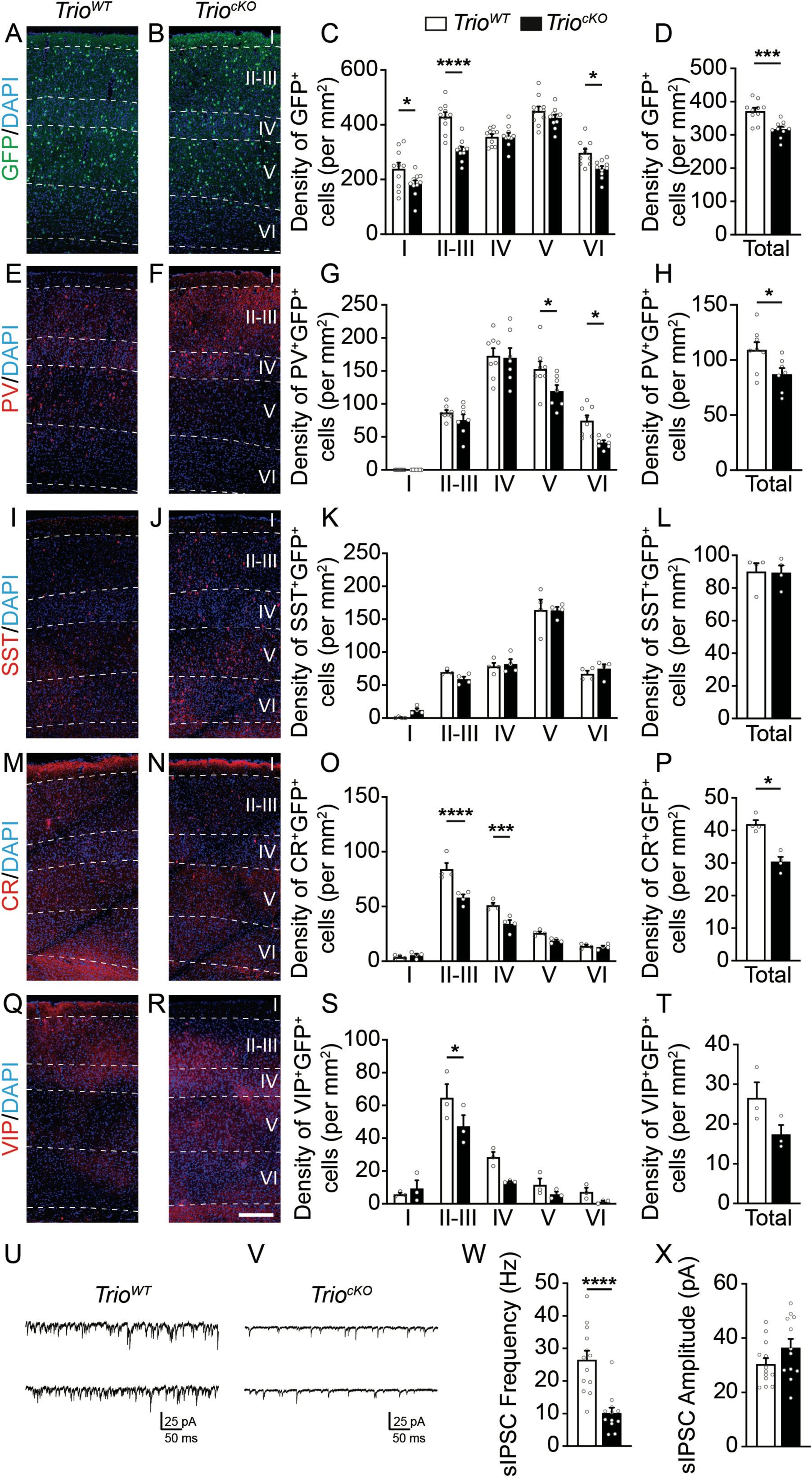
The targeted deletion of *Trio* in INs leads to reduced densities of cortical deep layer parvalbumin-expressing INs and of superficial layer calretinin and VIP-expressing INs in the post-natal somatosensory cortex, resulting in reduced cortical inhibition. (**A-D**) Reduced density of GABAergic INs in the somatosensory cortex of *Trio^cKO^* mice at P21. Examples of coronal brain sections showing fate-mapped GABAergic INs (GFP, green) and DAPI counterstaining (blue) at P21 in the somatosensory cortex of *Trio^WT^* (**A**) and *Trio^cKO^* (**B**) mice. (**C-D**) Histograms showing the quantifications of fate-mapped GABAergic INs at P21 in the mouse somatosensory cortex. (**E-T**) Reduced density of deep layer parvalbumin (PV)-expressing INs and superficial layer calretinin (CR)- and vasointestinal peptide (VIP)-expressing INs, but not somatostatin (SST)-expressing INs, in the somatosensory cortex of *Trio^cKO^* mice at P21. Immunofluorescence for PV (**E-F**), SST (**I-J**), CR (**M-N**) and VIP (**Q, R**) with DAPI counterstaining in the *Trio^WT^* (**E, I, M, Q**) and *Trio^cKO^* (**F, J, N, R**) somatosensory cortex at P21. Histograms of corresponding quantifications are shown for PV^+^GFP^+^ INs (**G-H**), SST^+^GFP^+^ INs (**K-L**), CR^+^GFP^+^ INs (**O-P**) and VIP^+^GFP^+^ INs (**S, T**) (n = 10 *Trio^WT^* and 10 *Trio^cKO^* brains for GFP quantifications; n = 8 *Trio^WT^* and 7 *Trio^cKO^* brains for PV quantifications; n = 4 *Trio^WT^* and 4 *Trio^cKO^* brains for SST and CR quantifications and n = 3 *Trio^WT^* and 3 *Trio^cKO^* brains for VIP quantification). (**U-X**) Reduced cortical inhibition in *Trio^cKO^* mice. Representative electrophysiological traces of sIPSCs in acute slices from *Trio^WT^* (**U**) and *Trio^cKO^* (**V**) mice. (**W-X**) Histograms showing the frequency (**W**) and amplitude (**X**) of sIPSCs (n = 13 cells from 4 *Trio^WT^* brains and 12 cells from 4 *Trio^cKO^* brains). \**P* < 0.05, \*\*\**P* < 0.001 and \*\*\*\**P* < 0.0001, by two-way ANOVA followed by Bonferroni’s multiple comparisons test (**C, G, K, O, S**) or by Mann-Whitney test (**D, H, L, P, T, W, X**). Scale bar: 200 µ m (**N**).

To assess whether this reduction of IN density results in functional alterations of cortical inhibition, we recorded spontaneous inhibitory post-synaptic currents (sIPSCs) in S1 layer V pyramidal cells in acute slices of P30 animals (Figure 2 U-X). We found a striking reduction in the frequency of sIPSCs in *Trio^cKO^* animals compared to *Trio^WT^* littermates (Figure 2 W), while the amplitude of sIPSCs remained similar between *Trio^WT^* and *Trio^cKO^* slices (Figure 2 X), compatible with a reduction of cortical IN numbers in adult *Trio^cKO^* mice.

### The conditional deletion of Trio impairs the tangential migration of INs and results in a premature switch to radial migration, with an accelerated entry in the cortical plate

To investigate whether prenatal alterations in IN migration contribute to this relative loss of INs in juvenile *Trio^cKO^* mice, we conducted cellular quantifications of fate-mapped MGE-derived cells at various embryonic stages. We found that, as in *Trio^-/-^* null embryos, IN migration is significantly altered in *Trio^cKO^* embryos compared to *Trio^WT^* littermates (Figure 3). At e13.5, corresponding to the peak migration of MGE-derived INs (including PV- and SST-expressing INs), INs migrate more slowly, with a 48 % (bin 4) to 96% (bin 5) reduction of INs reaching the medial dorsal pallium (bins 4-5) at this stage (Figure 3 A-C). We observe a premature switch towards a radial migration mode and a premature entry in the cortical plate of a subset of INs at e13.5 (Figure 3 D-E), although the majority (>80%) of INs remain in tangential migration at this time-point, even amongst those which have entered the cortical plate. Notably, we observed similar migration delays at e15.5 (Figure 3 F-K), corresponding to the peak migration age for CGE-derived INs (including CR- and VIP-expressing INs). We find a similar premature switch to radial migration of INs (Figure 3 K-L), together with a delayed migration of the still-tangentially migrating INs that fail to reach the medial dorsal pallium at this stage (Figure 3 O). Together, these data suggest that, in *Trio^cKO^*, tangentially migrating INs migrate more slowly than INs of *Trio^WT^* mice and that mutant INs fail to properly distribute across the dorsal pallium, resulting in a persistent reduction of cortical IN density at P21.

**Figure 3.**
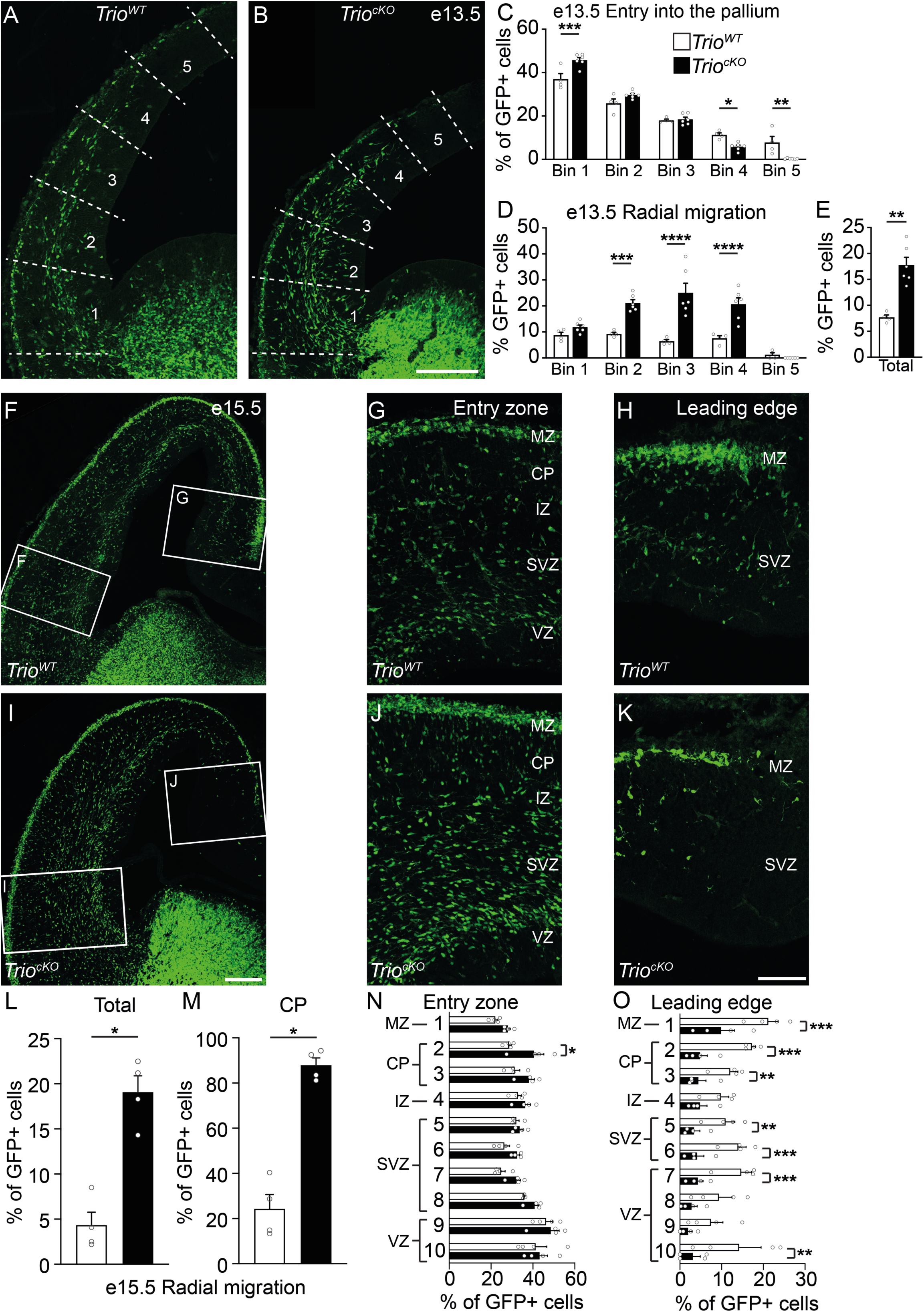
Conditional deletion of *Trio* in GABAergic INs delays tangential migration. (**A-E**) GABAergic IN migration at e13.5 is delayed in *Trio^cKO^* brains. Examples of coronal brain sections from e13.5 mouse embryos are shown for *Dlx5/6^Cre^;RCE^EGFP^* (*Trio^WT^*) (**A**) and *Dlx5/6^Cre^;Trio^c/c^;RCE^EGFP^* (*Trio^cKO^*) littermates (**B**), with binning strategy delineated by the dotted lines. (**C**) Histogram showing the distribution of GFP+ INs in the dorsal pallium of e13.5 *Trio^WT^* and *Trio^cKO^* brains. (**D-E**) Histograms showing the proportion of radially migrating GFP+ cells in the dorsal pallium at e13.5 (n = 4 *Trio^WT^* and 6 *Trio^cKO^* brains). (**F-O**) *Trio* deletion in GABAergic INs delays tangential migration at e15.5 and induces a premature switch to radial migration mode. Examples of coronal brain sections from *Trio^WT^* (**F**) and *Trio^cKO^* (**I**) littermates at e15.5. High magnifications of the entry zone (**G, J**) and the leading edge (**H, K**) are shown with the delimitation between the developing neocortical layers. (**L-M**) Histograms showing the proportion of radially migrating GFP+ INs in the entire neocortex (**L**) and in the cortical plate (CP) only (**M**). (**N-O**) Histograms showing the proportion of GFP+ INs at e15.5 across the dorsal pallium in the entry zone (**N**) and the leading edge (**O**) between *Trio^WT^* and *Trio^cKO^* brains. (n = 4 *Trio^WT^* and 3 *Trio^cKO^* brains). \**P* < 0.05, \*\**P* < 0.01, \*\*\**P* < 0.001, \*\*\*\**P* < 0.0001, by Mann-Whitney (**E, L, M**) or two-way ANOVA followed by Bonferroni’s multiple comparisons test (**C, D, N, O**). Scale bars: 200 µ m (**B** and **I**) and 50 µ m (**K**).

### Trio regulates structural remodeling and the dynamics of IN tangential migration

To investigate the cellular mechanisms underlying the alterations of IN migration in *Trio^cKO^* mutants, we next analyzed the morphology of developing INs in e13.5 *Trio^cKO^* embryos and *Trio^WT^* littermates using 3-D neuronal reconstitutions of cortical fate-mapped GFP-expressing GABAergic INs on fixed cryosections (Figure 4 A-B). We found that, in *Trio^cKO^* animals, migrating INs displayed a multipolar morphology, with more numerous leading and trailing neurites, and a larger cell body (Figure 4 C, D, G). Further, *Trio^cKO^* INs displayed excessively long and complex leading neurites and more complex trailing neurites compared to INs of *Trio^WT^* mice (Figure 4 E-F, H-I).

**Figure 4.**
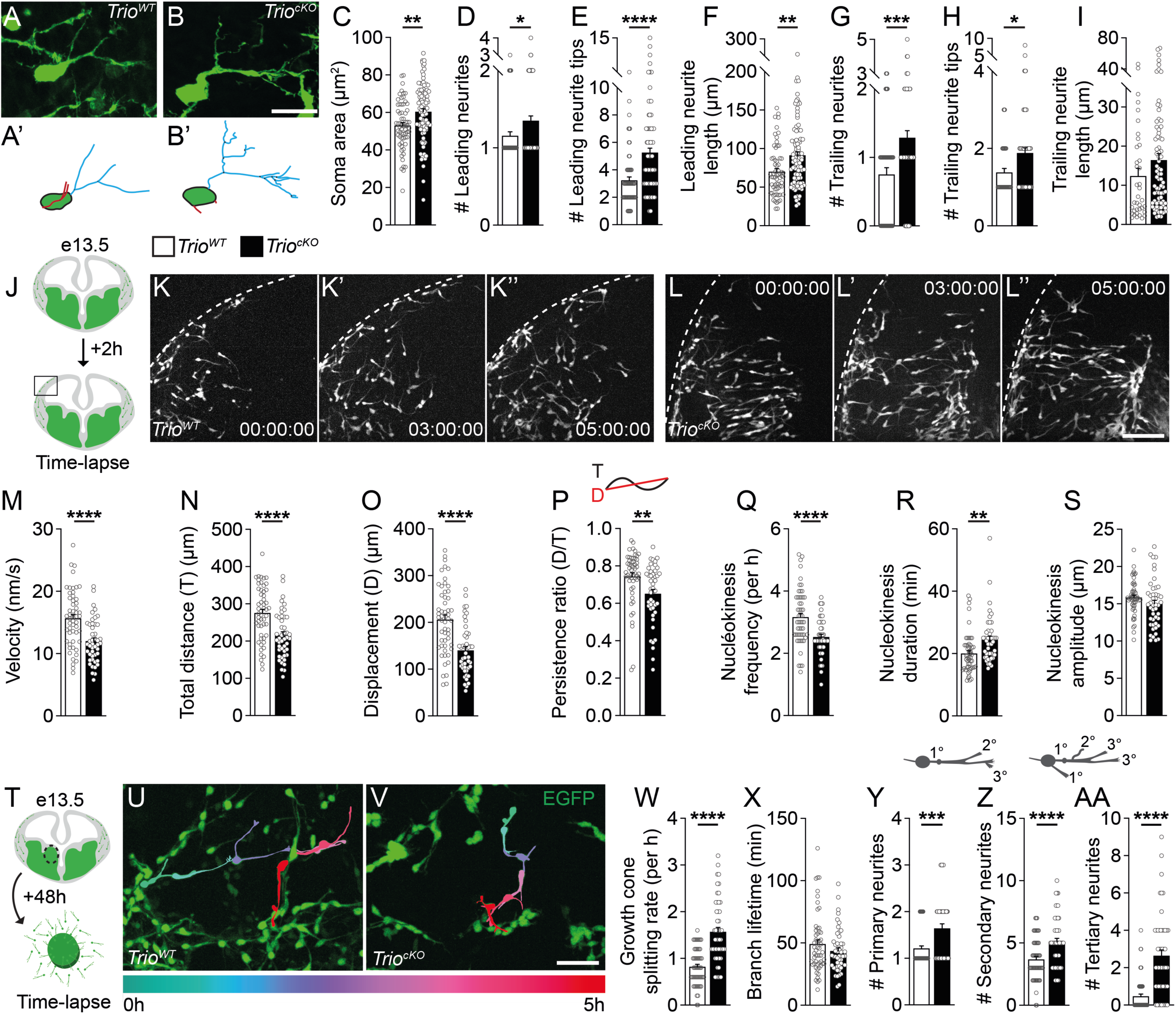
Conditional deletion of *Trio* in GABAergic INs induces morphological deficits in developing INs and impairs the dynamics of tangential migration. (**A-I**) The morphology of GABAergic INs at e13.5 is impaired in *Trio^cKO^* brains. Examples of GFP+ fate-mapped cortical INs imaged at the migratory front in 50 µ m-thick coronal brain sections from *Trio^WT^* (**A**) and *Trio^cKO^* (**B**) littermates at e13.5, with their respective 3D reconstitution (**A’**, **B’**). (**C-I**) Histograms showing measurements for (**C**) cell body area, (**D**) number, (**E**) complexity and (**F**) length of leading neurites, as well as (**G**) number, (**H**) complexity and (**I**) length of trailing neurites. (n = 64 cells from 3 *Trio^WT^* brains and 94 cells from 4 *Trio^cKO^* brains). (**J-S**) GABAergic IN migration and branching dynamics are impaired in *Trio^cKO^* embryonic brains. (**J**) Schematic of the experimental design for live-imaging of e13.5 organotypic brain slices after 2h in culture. Time-lapse sequences showing the migration of GFP+ INs in organotypic brain slices from *Trio^WT^* (**K-K’’**) and *Trio^cKO^* (**L-L’’**) mouse embryos. (**M-S**) Histograms showing (**M**) the migration speed, (**N**) the total distance, (**O**) the net displacement, (**P**) the directionality, as well as (**Q**) nucleokinesis frequency, (**R**) duration and (**S**) amplitude of *Trio^WT^* and *Trio^cKO^* GFP+ INs (n = 53 cells from 3 *Trio^WT^* brains and 46 cells from 3 *Trio^cKO^* brains). (**T**) Schematic of the experimental design for the live-imaging of MGE explant cultures. Time-lapse sequences showing the migration and branching of (**U**) GFP+ *Trio^WT^* and (**V**) *Trio^cKO^* INs in MGE explant cultures. (**W**) Histograms showing the frequency of growth cone splitting and (**X**) the lifetime of newly formed branches during migration. (**Y-AA**) Histograms showing the branch order classification during migration (n = 57 GFP+ MGE-INs from 3 *Trio^WT^* brains and 50 GFP+ MGE-INs from 3 *Trio^cKO^* brains). See also Suppl. Fig. 3 for data on migration dynamics and nucleokinesis in explants. \**P* < 0.05, \*\**P* < 0.01 and \*\*\*\**P* < 0.0001, by Student’s t-test. Scale bars: 20 µ m (**A**), 100 µ m (**L’’**) and 50 µ m (**V**).

To further investigate the impacts of the loss of *Trio* on IN migration dynamics, we conducted time-lapse imaging of fate-mapped GFP-expressing cells in acute organotypic brain slices of *Trio^cKO^* and *Trio^WT^* e13.5 embryos (Figure 4 J-L’’ and Supplementary videos 1-2). We observed that the migration dynamics of tangentially-migrating *Trio^cKO^* INs were significantly impaired compared to those of *Trio^WT^* INs. Indeed, we find a significant reduction in the mean velocity (Figure 4 M), the total distance travelled (Figure 4 N), the net displacement (Figure 4 O), the directionality (i.e. increased meandering) (Figure 4 P), and the frequency of nuclear translocations (Figure 4 Q) of *Trio^cKO^* INs compared to *Trio^WT^* INs. Nucleokinesis cycles in *Trio^cKO^* INs were of longer duration (Figure 4 R) but their amplitude was unchanged compared to those of *Trio^WT^* INs (Figure 4 S).

We next examined branching dynamics in migrating INs of *Trio^cKO^* and *Trio^WT^* embryos. Imaging of INs in organotypic slices does not provide sufficient resolution to analyze branching dynamics, given the tendency of INs to move out of the field of view. Therefore, we used MGE explants from *Trio^WT^* and *Trio^cKO^* e13.5 brains, maintained in culture for 48h before live-cell imaging (Figure 4 T-V and Supplementary videos 3-4). First, we observed deficits in the migration dynamics of *Trio^cKO^* INs migrating out of MGE explants compared to *Trio^WT^* INs (Supplementary figure 3 A-F), akin to what we had observed in *Trio^cKO^* INs migrating in acute slices. In addition, we observed significant alterations in branching dynamics in *Trio^cKO^* INs, with an increased growth cone splitting rate (Figure 4 W), but preserved mean lifetime of transient branches (Figure 4 X), together leading to a net increase in persistent new branches. Further, we observed an increased number of primary, secondary and tertiary neurites (Figure 4 Y-AA), consistent with the morphological deficits observed in fixed e13.5 *Trio^cKO^* brains. Altogether, our results suggest that *Trio* is a critical regulator of IN morphology and migration, with significant roles in the control of IN nucleokinesis and branching dynamics.

### TrioGEFD2-mediated RhoA activation regulates the morphological development of INs

The active branching and remodeling of the leading process in response to environmental cues helps guide the migration of INs ^29, 30^. This process involves the activation of Rho-GTPases at the tip of the leading process, including RhoA, Rac1 and Cdc42, by GEF proteins such as Trio ^30–33^. To dissect the contribution of each GEF domain to the processes involved in IN morphological remodeling and migration, we conducted a series of *ex utero* rescue experiments in *Trio^cKO^* mouse embryos by electroporating a control *Dlx5/6::mCherry* plasmid together with (1) the full-length *Trio* cDNA (*Trio^F-L^*); (2) a mutated version of *Trio* cDNA lacking the GEF1 domain, but with an intact GEF2 domain presumably able to activate RhoA (*Trio*^Δ*GEFD*^^1^); or (3) a mutated version of *Trio* cDNA lacking the GEF2 domain, but with an intact GEF1 domain presumably able to activate Rac1/RhoG/Cdc42 (*Trio*^Δ*GEFD*^^2^), followed by MGE explant cultures (Figure 5 A-E’). The explants were cultured for 48h, imaged with high-resolution time-lapse imaging for 5h and fixed and processed for 3D reconstitutions.

**Figure 5.**
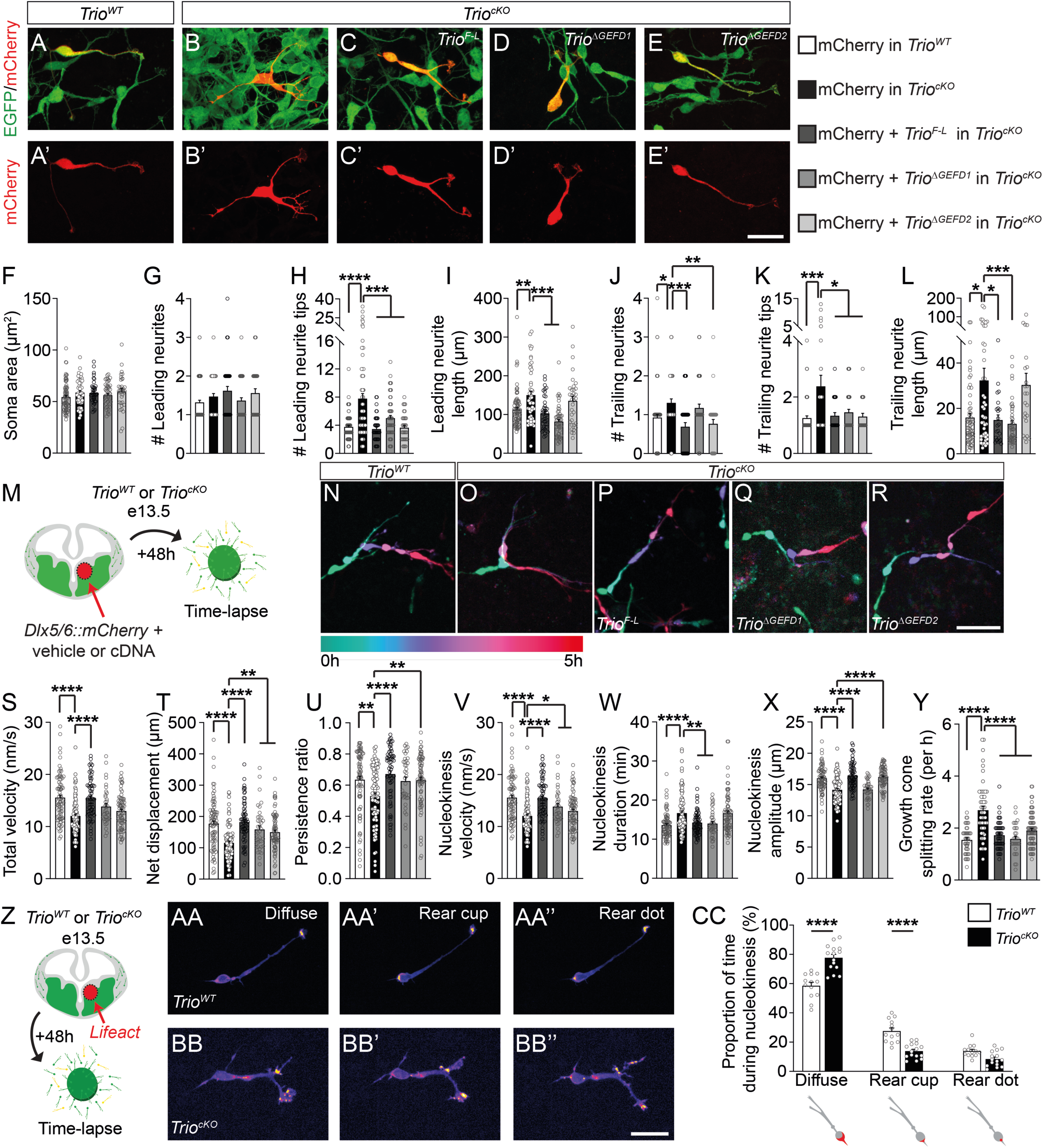
Both Trio-GEF domains are required for IN migration, and nucleokinesis dynamics defects are partly attributable to disruptions of actin remodeling. (**A-E’**) Confocal images of (**A-A’**) *Trio^WT^* INs in MGE explants cultures electroporated with a control *Dlx5/6::mCherry* construct (mCherry), (**B-B’**) *Trio^cKO^* INs in MGE explant cultures electroporated with mCherry or *Trio^cKO^* INs in MGE explant cultures electroporated with mCherry together with (**C-C’**) full-length *Trio* cDNA (*Trio^F-L^*), (**D-D’**) a mutated version of the *Trio* cDNA lacking the GEFD1 (*Trio*^Δ^*^GEFD^*^1^) or (**E-E’**) a mutated version of the *Trio* cDNA lacking the GEFD2 (*Trio*^Δ^*^GEFD^*^2^). (**F-L**) Histograms of (**F**) the soma area, (**G**) the number, (**H**) complexity and (**I**) length of leading neurites, as well as (**J**) the number, (**K**) complexity and (**L**) length of trailing neurites, following the rescue by *Trio^F-L^*, *Trio*^Δ^*^GEFD^*^1^ or *Trio*^Δ^*^GEFD^*^2^ cDNA (n = 82 GFP+ MGE-INs from 5 *Trio^WT^* brains electroporated with mCherry, 49 GFP+ MGE INs from 4 *Trio^cKO^* brains electroporated with mCherry, 42 GFP+ MGE-INs from 3 *Trio^cKO^* brains co-electroporated with mCherry and *Trio^F-L^*, 52 GFP+ MGE-INs from 6 *Trio^cKO^* brains co-electroporated with mCherry and *Trio*^Δ^*^GEFD1^* and 34 GFP+ MGE-INs from 4 *Trio^cKO^* brains co-electroporated with mCherry and *Trio*^Δ^*^GEFD2^*). (**M-Y**) Both GEF domains are required to fully rescue IN migration. (**M**) Schematic of the experimental procedure to rescue IN migration and branching. (**N**) Time-lapse color-coded sequences showing a *Trio^WT^* GFP+ MGE-IN co-electroporated with mCherry and vehicle, (**O**) a *Trio^cKO^* GFP+MGE-IN co-electroporated with mCherry and vehicle, (**P**) a *Trio^cKO^* GFP+ MGE-IN co-electroporated with mCherry and *Trio^F-L^*, (**Q**) a *Trio^cKO^* GFP+ MGE-IN co-electroporated with mCherry and *Trio*^Δ^*^GEFD1^* and (**R**) a *Trio^cKO^* GFP+ MGE-IN co-electroporated with mCherry and *Trio*^Δ^*^GEFD2^*. (**S-Y**) Histograms showing the rescue of (**S**) total migration speed (velocity), (**T**) net displacement (**U**) persistence ratio (directionality), (**V**) nucleokinesis cycle speed (**W**) nucleokinesis cycle duration and (**X**) nucleokinesis cycle amplitude (n = 81 GFP+ MGE-INs from 5 *Trio^WT^* brains electroporated with mCherry, 88 GFP+ MGE-INs from 6 *Trio^cKO^* INs electroporated with mCherry, 65 GFP+ MGE-INs from 3 *Trio^cKO^* brains co-electroporated with mCherry and *Trio^F-L^*, 38 GFP+ MGE-INs from 3 *Trio^cKO^* brains co-electroporated with mCherry and *Trio*^Δ^*^GEFD1^* and 76 GFP+ MGE-INs from 4 *Trio^cKO^* brains co-electroporated with mCherry and *Trio*^Δ^*^GEFD2^*). (**Y**) Histogram showing the rescue of the growth cone splitting rate (n = 42 GFP+ MGE-INs from 3 *Trio^WT^* brains electroporated with mCherry, 51 GFP+ MGE-INs from 4 *Trio^cKO^* brains electroporated with mCherry, 52 GFP+ MGE-INs from 3 *Trio^cKO^* brains co-electroporated with mCherry and *Trio^F-L^*, 29 GFP+ MGE-INs from 3 *Trio^cKO^* brains co-electroporated with mCherry and *Trio*^Δ^*^GEFD1^* and 69 GFP+ MGE-INs from 4 *Trio^cKO^* brains co-electroporated with mCherry and *Trio*^Δ^*^GEFD2^*. (**Z-CC**) Conditional deletion of *Trio* in GABAergic INs reduces actin remodeling. (**Z**) Schematic of the experimental design used to live-track actin dynamics in MGE-INs. (**AA-BB’’**) Time-lapse sequences showing the distribution of F-actin (fire mode) in a *Trio^WT^* (**AA-AA’’**) and a *Trio^cKO^* (**BB-BB’’**) GFP+ MGE-IN after electroporation of the Lifeact construct. (**CC**) Histogram showing the distribution of F-actin in the soma of GFP+ MGE-INs (n = 13 GFP+ MGE-INs from 3 *Trio^WT^* brains and 15 GFP+ MGE-INs from 4 *Trio^cKO^* brains). \**P* < 0.05, \*\**P* < 0.01, \*\*\**P* < 0.001 and \*\*\*\**P* < 0.0001 by one-way ANOVA followed by Tukey’s multiple comparisons tests (**F-L**, **S-Y**), or two-way ANOVA followed by Bonferroni’s multiple comparisons test (**CC**). Note that, with the exception of **S** and **V**, whenever *Trio^cKO^* MGE-INs electroporated with *Trio^F-L^*, *Trio*^Δ^*^GEFD1^* or *Trio*^Δ^*^GEFD2^* were statistically different from *Trio^cKO^* MGE-INs electroporated with mCherry (statistical differences currently showed on the figure), they were also not different from *Trio^WT^* MGE-Ins electroporated with mCherry. Thus, for better clarity, only comparisons with *Trio^cKO^* INs electroporated with mCherry (black column) are shown in **F-L** and **S-Y**, but *p*-values for all comparisons can be found in Supplementary Table 1. Scale bars: 25 µ m (**E, BB’’**) and 50 µ m (**R**).

On the fixed explants, we confirmed that, as before, *Trio^cKO^* INs electroporated with the control *Dlx5/6::mCherry* plasmid displayed significant morphological deficits compared to electroporated INs of *Trio^WT^* mice; with longer and more complex leading and trailing neurites, along with an increased number of trailing neurites (Figure 5 F-L). Further, the co-electroporation of the *Dlx5/6::mCherry* plasmid with the *Trio^F-L^* construct in *Trio^cKO^* INs rescued these morphological deficits (Figure 5 F-L), confirming that *Trio* loss-of-function in INs was indeed responsible for the morphological deficits reported above. Interestingly, the expression of the *Trio*^Δ*GEFD*^^1^ cDNA in *Trio^cKO^* INs also rescued most of the morphological deficits, including the increased leading and trailing neurite complexity and length, suggesting an important role for the RhoA-activating GEFD2 domain in these processes. Conversely, the expression of the *Trio*^Δ*GEFD*^^2^ cDNA rescued the complexity of the leading and trailing neurites, as well as the number of trailing neurites, but not the leading and trailing neurite length (Figure 5 F-L). Thus, our data suggest a requirement of the GEFD2-RhoA pathway for the proper regulation of leading and trailing process morphology in tangentially migrating INs, with a redundant contribution from the GEFD1-Rac1/Cdc42 pathway in controlling leading and trailing neurite complexity.

### Trio GEFD1 and GEFD2 domains redundantly control leading neurite branching dynamics, but both are required for proper cortical IN migration

We next analyzed the requirement of Trio-GEFD1 and Trio-GEFD2 in rescuing the dynamics of neurite remodeling and migration by analyzing time-lapse imaging of the electroporated MGE explants described above (Figure 5 M-R and Supplementary videos 5-7). As before, *Trio^cKO^* INs electroporated with the control *Dlx5/6::mCherry* plasmid displayed several deficits in migration dynamics, when compared to *Trio^WT^* INs, as well as impairments in the leading process branching dynamics (Figure 5 S-Y and Supplementary Figure 3 H-O). As expected, the expression of the *Trio^F-L^* plasmid in *Trio^cKO^* INs rescued most aspects of the migration and branching dynamics deficits (Figure 5 S-Y and Supplementary Figure 3 H-O).

When investigating the contribution of each GEF domain to the migration and branching dynamics, we observed that deficits in the duration of nucleokinesis cycles (Figure 5 W) and in the number of primary neurites (Supplementary Figure 3 M) induced by *Trio* conditional deletion were exclusively rescued by the electroporation of the *Trio*^Δ*GEFD*^^1^, whereas impairments in migration directionality (Figure 5 U) and nucleokinesis amplitude (Figure 5 X) were exclusively rescued by the *Trio*^Δ*GEFD*^^2^. As a result, electroporation of either *Trio*^Δ*GEFD*^^1^ or *Trio*^Δ*GEFD*^^2^ rescued the net soma displacement (Figure 5 T), as well as the frequency of growth cone splitting (Figure 5 Y) and the number of third order neurites (Supplementary Figure 3 O), and partially rescued nucleokinesis cycle velocity (Figure 5 V and supplementary table 1), suggesting a redundancy in controlling these specific dynamic processes. However, neither construct was able to rescue the global migration velocity (Figure 5 S), the total distance (Supplementary Figure 3 H) nor the nucleokinesis cycle frequency by itself (Supplementary Figure 3 I). Altogether, these data suggest that the two GEF domains exert a complementary control over different aspects of migration and branching dynamics, with some redundancy in terms of net displacement, nucleokinesis velocity and growth cone splitting. Nonetheless, both domains are required for the proper net migration of GABAergic INs (Figure 6).

**Figure 6.**
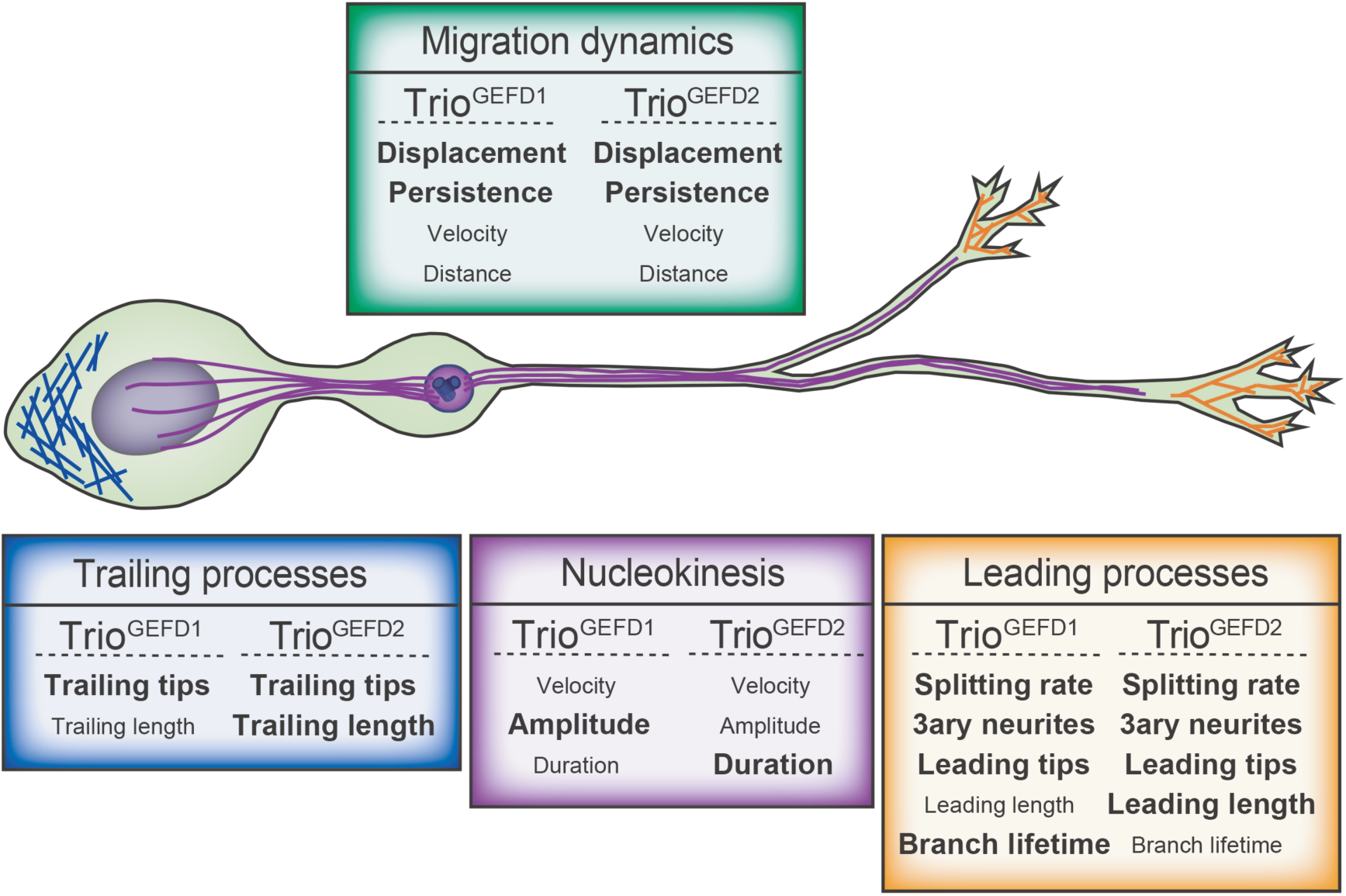
Summary schematic illustrating the contribution of GEFD1 and GEFD2 in regulating GABAergic IN migration dynamics and morphology. At the molecular level, both GEFD1 and GEFD2 contribute to the tangential migration of GABAergic INs. Trio GEFD1 regulates the amplitude of nucleokinesis and the migration directionality, while Trio GEFD2 controls nucleokinesis cycle duration (purple box; rescued parameters in bold). Thus, deleting either domain results in a significant delay of IN migration, and both domains are required to fully rescue the net velocity and distance traveled by INs in *Trio^cKO^* mice (green box; rescued parameters in bold). Further, both domains independently regulate leading process branching dynamics (orange box; rescued parameter in bold), but Trio GEFD2 is critical in restricting neurite length and is thus required to fully rescue IN morphology in *Trio^cKO^* mice (blue box and orange box; rescued parameters in bold).

### Trio controls IN migration by regulating actin cytoskeleton remodeling

Nucleokinesis, which involves a translocation of the nucleus in the leading process, is regulated by actin remodeling and actomyosin contractions at the rear of the cell body ^29, 34–36^. Thus, we investigated the impact of *Trio* deletion on actin remodeling in GABAergic INs by generating MGE explants from *Trio^WT^* and *Trio^cKO^* e13.5 embryos after *ex utero* MGE electroporation of the Lifeact plasmid (Figure 5 Z). During IN migration, F-actin can be found in the cell body either in a non-polarized diffuse state, forming a cup at the rear of the cell body in preparation for actomyosin contractions, or condensed in a dot at the rear of the cell body, just before nuclear translocation (Figure 5 AA-BB) ^36, 37^. In migrating *Trio^cKO^* INs, F-actin distribution remains diffuse, largely failing to form a rear cup, when compared to *Trio^WT^* INs (Figure 5 CC, Supplementary videos 8-9). Together, these data suggest that *Trio* controls actin turnover as well as its localization in order to promote the actomyosin contractions required for successful and rapid nucleokinesis. Thus, mislocalization of polymerized actin and failed actomyosin contractions in *Trio^cKO^* INs likely contribute to the slower and inefficient nucleokinesis observed in *Trio^cKO^* INs, and likely reduces the speed and distance travelled by tangentially migrating INs in *Trio^cKO^* mice.

## Discussion

This study identifies the dual GEF Trio protein as a major regulator of the morphological development and migration of cortical GABAergic INs, suggesting that major features of *TRIO*-associated NDDs reflect a developmental interneuronopathy. In particular, we find that the deletion of *Trio* in GABAergic INs results in spontaneous seizures and ASD-like behaviors in *Trio^cKO^* mutant mice. This phenotype is largely attributable to a delayed migration of INs, resulting in reduced PV+ and VIP/CR+ IN numbers in the mature cortex, together with decreased cortical inhibition in mutant mice. Mechanistically, we find that the deletion of *Trio* alters the branching and nucleokinesis dynamics during IN tangential migration. While Trio GEFD1 and GEFD2 independently regulate specific steps of IN migration, both are required to fully rescue the migration dynamics. Notably, we observe disruptions of actin remodeling, with deficits in actin polymerization and actomyosin contractions, slowing nucleokinesis in *Trio^cKO^* INs. Altogether, our findings suggest that *TRIO*-associated NDDs are developmental interneuronopathies, with both GEF domains playing critical roles during IN migration.

### Interneuronopathies in NDDs

Human *de novo* and recessive *TRIO* mutations result in a spectrum of NDDs, including early-onset developmental epileptic encephalopathies (DEE) and/or ASD with or without ID ^1–6^. The mechanisms underlying these clinical phenotypes are not fully understood. Our results suggest that the loss of *Trio* is particularly detrimental to the tangential migration of GABAergic INs, as also suggested by others ^16^, and that its selective deletion in INs suffices to impair cortical inhibition and to induce epilepsy and ASD-like phenotypes in mice. This finding expands a growing body of evidence suggesting that a primary deficit of cortical inhibition, and particularly of PV-expressing INs (PV-INs), results in epilepsy and ASD-like behaviors in rodents ^38–42^. PV-INs are the main inhibitory targets of thalamocortical afferents and they provide strong and rapid synaptic inhibition on the soma of adjacent pyramidal cells, therefore regulating cortical excitability ^43–49^. Thus, the deletion of genes involved in the specification ^50^, proliferation^51^, migration and maturation ^26, 39, 52–56^, excitability ^57–60^ or synaptic function ^40, 41, 61^ of PV-INs result in epilepsy and cognitive deficits in mice. Furthermore, PV-INs participate in the generation of gamma oscillations, known to be critical for adequate cortical processing ^62, 63^, attention ^64^ and memory ^65^. Disruption of gamma oscillations may thus contribute to the array of cognitive and behavioral deficits associated with PV-INs impairments ^38, 42, 56, 61, 66^, including some of the deficits observed in *Trio^cK^* mice.

Importantly, deficits in other IN subtypes also likely contribute to the phenotype observed in *Trio^cKO^* mice. CR/VIP-expressing INs orchestrate cortical synchrony through local and long-range disinhibition ^67^ and dysfunction of VIP-INs results in ASD/DEE phenotypes in rodents ^68, 69^. Thus, the reduction of superficial cortical layer CR and VIP-expressing INs in *Trio^cKO^* mice might contribute to their overall neurobehavioral phenotype.

Surprisingly, our quantification results contradict the findings reported by Sun, Wang ^16^ who did not observe a reduction of IN densities in the somatosensory cortex of *Dlx5/6^Cre^;Trio^c/c^* mice. Methodological differences and increased statistical power in our study might contribute to our observations of differences in IN densities. Further, Sun, Wang ^16^ did not observe spontaneous seizures but found a reduced threshold to PTZ-induced seizures in *Dlx5/6^Cre^;Trio^c/c^* mice. This difference may stem from age differences in the study populations. Indeed, our study was conducted in early adult mice (P21-P30), while Sun et al. investigated older adult mice (P35-P60). A transient developmental window for seizure susceptibility occurs between P20-P25 in various models of PV-IN dysfunction, with reduced seizure susceptibility at later stages (P33-45) ^40, 41, 61, 70–72^. By contrast, cognitive and behavioral deficits persist in adult ages ^61, 73^, suggesting that early disruption of PV-INs development results in long-lasting impacts on cognition and behavioral control.

### Impaired tangential migration as a disease mechanism for Trio-associated NDD

Trio is a key regulator of the development ^1, 7, 14^, migration ^13, 15^ and synaptic function of excitatory neurons ^4, 7, 19^. Our findings and those of Sun, Wang ^16^ suggest that Trio is also a critical regulator of the morphological development and migration of GABAergic INs, in addition to many other well-described molecular actors (reviewed in ^32, 38^).

### Trio regulates leading process branching and prevents cortical plate invasion

The process of tangential migration involves a variety of intrinsic and extrinsic cues to attract migrating neurons in the proper direction, while guiding them away from non-cortical structures, maintaining INs in their proper migratory streams ^32, 35^. Our findings, aligned with those of Sun, Wang ^16^, reveal that, in the absence of *Trio*, tangentially migrating INs switch rapidly to a radial migration mode, leading to a premature entry in the cortical plate. This premature entry in the cortical plate reflects a loss of response to SDF1/CxCr4 signaling in *Trio* cKO INs ^16^. Indeed, SDF1/CxCr4 signaling typically repels INs from the cortical plate, in a GEFD1/Rac1-dependant manner ^16, 74, 75^, until they down-regulate the CxCr4 receptor Cxcr12 ^76, 77^. Nonetheless, this premature entry in the cortical plate affects only a small proportion of migrating INs at e13.5 and e15.5, with more than 80% of migrating INs remaining in tangential migration in *Trio^cKO^* mutants at these time points. This mechanism is thus unlikely to explain the extent of the deficits observed in *Trio^cKO^* mutants, suggesting that other mechanisms affecting the dynamic processes of migration itself are likely also involved.

Our data suggest that the loss of *Trio* significantly impairs the dynamics of leading process branching, growth cone splitting, trailing process retraction and nucleokinesis, all key determinants of the overall speed and integrity of IN migration. Importantly, we show that these processes require both the GEFD1-Rac1/RhoG/Cdc42 and the GEFD2-RhoA pathways, as neither domain is sufficient to fully rescue IN migration. Notably, IN morphology, and in particular the length of both the leading and trailing neurites, is regulated by the GEFD2-RhoA pathway. By contrast, the retraction of the trailing neurite, which contributes to maintaining a bipolar morphology, depends on the GEFD1-Rac1/Cdc42 pathway. However, we find that both GEFD1 and GEFD2 pathways regulate branching dynamics and neurite complexity. This is concordant with previous reports of Rac1 (and Rac3) controlling the polarity and neurite complexity in tangentially-migrating INs ^78, 79^. However, the contribution of the RhoA pathway at this stage in INs had not been shown before. Similar mechanisms have also been documented in pyramidal cells, where both Rac1-dependent activation of the PAK/LIMK pathway and RhoA-dependent activation of ROCK/LIMK prevent axonal outgrowth ^79, 80^ and excessive neurite branching during radial migration ^79, 81–83^. Further, Cdc42 participates in the regulation of morphology and polarity in excitatory neurons ^33, 84^, while promoting migration of cerebellar granule cells through facilitation of radial-glia interactions ^84^ and axonal outgrowth in a TrioGEFD1-Cdc42-dependent (and Rac1 independent) manner ^14^. Whether Cdc42 plays a role in IN migration remains to be determined.

Interestingly, we find that both GEFD1 and GEFD2 pathways control nucleokinesis during tangential migration: GEFD2/RhoA determines the duration while GEFD1/Rac1 regulate the amplitude of each cycle. Although previous reports suggested that RhoGTPases, including RhoA, Rac1 and Cdc42, were dispensable during IN tangential migration but were required for progenitor cell division and migration competency early on, based on cell density assessment in fixed sections of KO mice ^81, 85–87^, our live-cell imaging data demonstrates a critical role of both GEFD1/Rac1 and GEFD2/RhoA pathways on the dynamic process of nucleokinesis in INs. This is aligned with other reports suggesting contributions of various Rho-GTPase regulators, including Trio, in promoting IN motility ^36, 37, 79, 88^. Further, we find that GEFD1/Rac1 helps maintain the directionality of migration in *Trio^cKO^* INs, a process known to depend on Rac1 activation^79^.

Interestingly, similar complementary roles have been described in migrating pyramidal cells, in which centrosome translocation initiating nucleokinesis requires Rac1 activation, in a spatially and temporally regulated fashion, via POSH signaling ^89^, while RhoA activation increases the rate of migration ^25, 90^ in a TrioGEFD2-dependent manner ^25^.

Further, in hindbrain pre-cerebellar neurons, RhoA regulates nuclear translocations, while Rac1 and Cdc42 are required for the initial extension of the leading process during tangential migration ^91^. Thus, the tight spatiotemporal regulation of multiple RhoGTPases by Trio appears to control the dynamics of migration of multiple neuronal cell types, including INs.

### Trio controls cytoskeletal actin remodeling during the tangential migration of INs

At the molecular level, RhoA, Rac1 and Cdc42 are known regulators of the cytoskeleton, controlling the remodeling of actin filaments and microtubules in response to various cues during neuronal migration (see review: ^33^. For instance, during tangential migration, the branching and elongation of the leading process occurs through stabilization of actin filaments by Filamin A ^92^. Notably, Filamin A binds Trio at the PH1-GEF1 domain, enabling its translocation to the actin cytoskeleton ^93, 94^. We find that the loss of Trio induces excessive elongation and branching of the leading process, perhaps suggesting an enhanced availability of Filamin A in the absence of Trio, leading to enhanced actin filaments stabilization through Trio-independent mechanisms ^95^. In addition, reduced activation of Rac1 or Cdc42, but not RhoA, decreases actin polymerization at the leading edge and alters microtubule dynamics in migrating cells, leading to impaired neuronal migration, aberrant polarity and excessive neurite branching ^78, 82, 84, 88, 96, 97^. The excessive branching dynamic observed in *Trio^cKO^* INs may thus reflect decreased actin polymerization and enhanced microtubule dynamics in the neurites, partly through hypoactivation of Rac1. Notably, Rac1, Cdc42 and RhoA have cell-specific and locally restricted interdependent relationships, as they sometimes mutually inhibit each other although, in some situations, they can activate each other ^98–101^. Notably, the PH2 domain next to Trio-GEFD2 negatively regulates RhoA ^102^. Binding of G-coupled protein Gqα to the PH2 relieves this inhibition and leads to targeted RhoA activation in a spatially restricted fashion ^102^. Further, Trio itself is a molecular target of both RhoA and Cdc42, suggesting complex feedback regulatory mechanisms ^84, 103^. The loss of Trio may thus result in local overactivation of RhoA, further inhibiting proper Rac1 activation at the leading edge, accentuating the impact described above on actin and microtubule dynamics in neurites.

Cytoskeletal remodeling also plays a role in the regulation of nucleokinesis. Indeed, during nucleokinesis, actin condensation at the rear of the nucleus supports the actomyosin contractions that propel the nucleus forward in both pyramidal cells and INs^34^. This process requires local RhoA activation ^37, 88, 96^. Here, we find that Trio promotes actin polymerization and condensation at the rear of the cell body, supporting nucleokinesis, while its loss results in a failure of proper actin condensation. Further, the frequency of each cycle is GEFD2/RhoA-dependent, consistent with impaired targeted activation of RhoA at this site in the absence of Trio.

In addition, actomyosin contractions during nuclear translocation requires dynamic microtubule reorganization in the cell body ^34^, while microtubule stabilization is required for the forward motion of the nucleus towards the centrosome in migrating INs ^78^. Trio is known to associate with acentrosomic microtubules to control cell polarity during migration ^104^. It also binds microtubule plus ends of growing microtubules during neurite extension ^105^. Thus, altered microtubule dynamics likely also contributes to some aspects of IN migration deficits observed in *Trio^cKO^* mutants.

In summary, our findings suggest that the DEE/ASD/ID-associated RhoGEF Trio regulates the development and migration of GABAergic INs by controlling cytoskeletal remodeling critical for proper motion of tangentially-migrating INs. Importantly, we demonstrate that while each GEF domain individually contributes to fundamental aspects of IN migration, with critical contributions from the GEFD2-RhoA pathway, their synergistic effects are required to ensure proper IN migration. These findings support an emergent literature suggesting that a subset of genetically determined DEE/ASD/NDD result from a primary impairment of IN migration (see reviews ^38, 106, 107^, and that such disorders may ultimately benefit from pharmacological or cell-based therapies aiming to reestablish circuit inhibition ^108–111^.

## Acknowledgments

L.E. was funded through a Savoy Foundation postdoctoral fellowship, a Fonds de recherche du Québec-Santé (FRQ-S) postdoctoral fellowship, a Sainte-Justine Foundation fellowship, a postdoctoral fellowship from the Transforming Autism Care Consortium Quebec Autism Research Training Program and a Brain Canada Research Fund, with financial support from Health Canada and the Kids Brain Health Network. P. K. R. and X. J. received a Savoy Foundation postdoctoral fellowship. J.-C. L. was supported by a Research Center grant (Centre Interdisciplinaire de Recherche sur le Cerveau et l’Apprentissage; CIRCA), from the Fonds de recherche du Québec – Santé (FRQ-S), a Group grant from Université de Montréal (Groupe de recherche sur la signalisation et circuiterie neurale; GRSNC) and is the recipient of the Canada Research Chair in Cellular and Molecular Neurophysiology (CRC 950-231066). E. R. was awarded FRQ-S and CIHR career awards and is the current holder of a Tier II Canada Research Chair on the Neurobiology of Epilepsy. L. L. and E. B-G. are INSERM investigators. This project was funded through operating grant and awards from CURE Epilepsy, the Savoy Foundation, the Scottish Rite Foundation, the Epilepsy Canada Foundation, the Fonds de recherche du Québec en Santé (FRQ-S) and the Canadian Institutes for Health Research (CIHR).

We would like to thank Sonia Garel for stimulating discussions, James Waldron for his technical support, the summer interns who participated in this project (Felicia Hansson, Gabrielle Godin, Petra Tamer, Lucas Toussaint, Louis Fauteux-Loiselle, Marc-André Ranger and Estelle Douet), Elke Küster-Schöck for her technical support at the Plateforme d’Imagerie Microscopique (PIM) of the CHU Sainte-Justine Research Center, as well as Denise Carrier and the animal care workers at the CHU Sainte-Justine Research Center.

## Conflict of interest

The authors declare that they have no conflict of interest.

## Material and methods

### Animals

All experiments were conducted in accordance with the Canadian Council of Animal Care (CCAC) guidelines, or with French and European regulations, and approved by the Université de Montréal Animal Care Committee and the CHU Sainte-Justine Animal Ethics Board, or by the Paris Descartes (CEEA 34) ethical committee for animal experimentation, in accordance with the information provided by the French Ministry of Research. *Dlx5/6^Cre^;Trio^c/c^;RCE^EGFP^* mutant mice (*Trio^cKO^*) were obtained by breeding *Dlx5/6^Cre^* mice ^26, 27^ with conditional *Trio^c/c^* mice ^13^ expressing the Cre-reporter allele *RCE^EGFP^* ^28^ to genetically label GABAergic INs. *Dlx5/6^Cre^;RCE^EGFP^* littermates (*Trio^WT^*) were used as controls throughout this study and conditional mutant mice are maintained on a *129sv* genetic background. However, for electrophysiology experiments and behavioral tests only, and to optimize the use of each litter, *Trio^c/c^*, *Trio^c/+^* (i.e. without Cre recombination) and *Dlx5/6^Cre^* littermates (i.e. with WT alleles) were all pooled as control animals and were compared to *Dlx5/6^Cre^;Trio^c/c^* conditional mutants. *Trio^-/-^* embryos were obtained by crossing *Trio^+/-^* mice ^17^ maintained on a BALB/c genetic background. PCR-based genotyping was performed as previously described ^13, 15^. Animals used to generate the conditional mutant mouse model were housed at the CHU Sainte-Justine animal facility under 12h light/dark cycles with water and food *ad libitum*. *Trio^+/-^* animals were housed and crossed at CDTA-TAAM Orléans. For pre-natal experiments, timed-pregnancies were monitored daily, with detection of a vaginal plug corresponding to embryonic stage e0.5.

### Video-EEG recordings

P30-36 mice were anesthetized with isoflurane (3% in O_2_, 0.8L/min) and bilateral stainless steel electrodes (Plastic One, E363/3/SPC) were implanted in the somatosensory cortex and CA1 areas (AP/ML/DV: Cortex: -1.0/±1.5/-1.0 mm, CA1: -2.0/±1.5/-2.0 mm to Bregma), with a reference electrode over the cerebellum ^40^. EEGs were recorded 24h later at 2000 Hz acquisition speed, filtered at HP 1.0 Hz and LP 70 Hz, and digitized using Cerevello-NeuroMed (Blackrock Microsystems). Recording sessions were conducted continuously for at least 72h in control and mutant littermates. Seizures were characterized as sustained epileptic activity > 4 seconds with behavioral changes and were quantified in terms of frequency, duration, type and severity on the modified Racine scale, as previously described ^40^.

### Patch-clamp electrophysiology

Brain slices of the somatosensory cortex (S1) were prepared as previously described ^40, 41, 61^ from control and mutant littermates at P30. Spontaneous inhibitory post-synaptic currents (sIPSCs) were recorded in layer V PCs in the presence of 10 µ M CNQX and 50 µ M DL-APV at -70 mV. Data were recorded and analyzed as before ^40, 41, 61^.

### Behavior

Behavioral tests were done at P30 in control and mutant littermates at a similar daytime period, by investigators blinded to the genotype. Mice were placed in the experimental room for at least 30 min and mice of both sexes were used in all experiments. All assays were conducted under video-tracking (Logitech c615 camera, SMART tracking system, Harvard Apparatus). The behavioral tasks were conducted sequentially from P30 onwards: Open Field (P30), Elevated Plus Maze (P32), Morris Water Maze (P34). Separate sets of mice underwent the Object Recognition, Marble Burying and Three Chamber Maze tasks. Except for the Marble Burying task, all behavioral assays were conducted and analyzed as in ^61^.

#### Open field

Mice were introduced at the center of the open-field arena (50 cm x 50 cm x 35 cm, length x width x height [L x W x H], illuminated with dim light) and movement and exploration were video-recorded for 10 min. The surface area was divided into center and periphery as 36% and 64% of the total area, respectively. To measure exploratory behavior, total distance travelled during the 10 minutes period, the speed and time spent in the center or the periphery and the number of entries in the center zone were calculated. The open-field arena was cleaned with 70% ethanol between each trial.

#### Elevated plus maze

Mice were introduced onto the center (5 x 5 cm) of an Elevated plus maze (EPM) facing a closed arm (arms 30 cm x 6 cm x 35 cm [L x W x H]), elevated 50 cm above the ground). The exploratory behavior was video-recorded for 5 min. The time spent and number of entries into open arms, closed arms and central zone were measured.

#### Novel object recognition

The novel object recognition test was performed in the above-mentioned open-field arena. After freely exploring the arena for 5 min, mice were recorded while exploring 2 similar small Falcon tissue culture flasks (50 ml volume) for 10 min and then returned to their home cages. After 30 min of inter-trial interval, one flask was replaced with a novel object with a different texture and duration of interaction was assessed in a second 10 min trial. Exploration time for each object was measured and represented as a discrimination index (time spent with novel object/total time spent with novel + familiar objects). Animals were scored as interacting with the objects when their nose was in contact with the object, or pointing toward the object within a defined distance (1.5 cm).

#### Morris Water Maze

Spatial memory was tested in a Morris water maze setup. A circular pool (125 cm) was filled with water and painted with non-toxic white paint (Brault & Bouthilier) and kept overnight at room temperature. Mice were placed in the pool and were expected to locate a submerged escape platform using the visible cues present in the room. Mice were trained for 5 consecutive days, 4 trials per day with a 1-3 min inter-session interval. In each trial, mice were first placed on the platform for 30 s, and then placed in the water at a random start position and allowed to swim to find the platform. Mice that were unable to find the platform within 60 s were placed back on the platform by hand. On day 6, a probe test was performed with the platform removed. Mice were placed in the pool opposite from the platform location and allowed to swim for 60 s. During training trials, the latency to reach the platform was measured and in the probe trial, the time spent in each quadrant of the pool was measured.

#### Marble burying task

Repetitive behavior was assessed using the marble burying task. Each mouse was placed in a clean cage with 3-inch bedding material and allowed to freely explore for 30 minutes. Then, 20 marbles were positioned equidistantly in the cage and each mouse was observed for an additional 20 minutes. The number of marbles buried during the 20-minute trial was quantified.

#### Three-chamber maze

Socialization assays were tested in the Three Chamber Maze, which consists of a rectangular arena of 70 x 45 cm surrounded by 30 cm tall transparent walls and separated in three equally-sized zones by two 45 x 30 cm transparent walls. The dividing walls had a door allowing the mice to circulate between zones. The animals were first placed in the middle of the central chamber and allowed to explore all empty chambers for 5 min. After habituation, an unfamiliar mouse of the same age and sex Stranger 1, S1) was placed inside a small wired cage and an empty cage was added in the opposite zone, while the middle zone remained empty. The tested animal was allowed to freely explore the three chambers of the arena for 10 min. Then, a new unfamiliar mouse of the same age and sex (Stranger 2, S2) was placed in the previously empty cage and the tested mouse was observed for an additional 10 min to assess preference for social novelty. S1 and S2 animals originated from different home cages and had never been in contact with the tested mice or between each other. The sociability and social novelty were quantified as the time spent by the tested animal in each chamber in close proximity with the other animal (or empty cage) during the first and second 10-min trial, respectively. The maze and cages were cleaned with 70% ethanol at the end of the task.

### Histology

Timed-pregnant females were sacrificed by cervical dislocation and embryos were collected at e13.5, e14.5, e15.5 or e18.5 as previously described ^15, 112^. Animals at P21 were deeply anesthetized with Pentobarbital (100 mg/kg, i.p.). Brains from e13.5 and e14.5 embryos were dissected out of the skull, rinsed in phosphate-buffered saline (PBS; 100 mM, pH 7.4) and fixed by immersion in 4% PFA overnight (ON) at 4°C. E15.5 and e18.5 embryos were perfused transcardially with 0.5 mL of 4% PFA and P21 animals were perfused transcardially with 12 mL of PBS followed by 12 mL of 4% PFA. E13.5, e14.5, e15.5, e18.5 and P21 brains were rapidly dissected out, post-fixed by immersion in 4% PFA for 1h (e13.5, e15.5 and P21) or ON (e14.5 and e18.5) at 4°C and cryoprotected in 30% sucrose in PBS. Then, brains were embedded in CryoMatrix and frozen on dry ice. Brains were cut along the coronal plane with a Cryostat (Leica) into 18 µ m-thick sections and collected on pre-cleaned Superfrost Plus Microscope Slides (Fisherbrand, Fisher Scientific, Cat. No. 12-550-15). Microscope slides were kept at -20°C until use. For e13.5 brains, every ninth section was cut at 50 µ m thickness, collected in PBS and stored at 4°C for further use (3D neuronal reconstitution experiments).

Fixed (e13.5, e15.5, P21) and free-floating (e13.5) sections were rinsed 3x10 min in PBS followed by blocking for 1h at room temperature (RT) in a solution containing 10% normal goat serum and 1% Triton X-100 in PBS. Sections were then incubated ON at 4°C in a blocking solution containing 5% normal goat serum and 0.1% Triton X-100 with the proper combination of primary antibodies: rat anti-EGFP (Cat. #04404-84, Nacalai Tesque; 1:1000), mouse anti-NeuN (Cat. #MAB377, Millipore; 1:1000), rabbit anti-parvalbumin (Cat. #PV-27, Swant; 1:1000), mouse anti-parvalbumin (Cat. #P3088, Sigma-Aldrich; 1 :1000), rabbit anti-somatostatin (Cat. #PA5-82678, ThermoFisher; 1:1000), rabbit anti-calretinin (Cat. #ABN2191, Millipore; 1:1000), rabbit anti-VIP (Cat. #20077, Immunostar; 1:1000). After several rinses in PBS, sections were then incubated for 2h at RT in the same blocking solution containing 1:1000 dilutions of corresponding Alexa secondary antibodies (Invitrogen) made in goat and coupled to either 488, 594 or 647 fluorophores. Sections were then rinsed several times in PBS before they were coverslipped with Vectashield® Hardset™ Antifade Mounting Medium (Cat. #H-1400, Vector Laboratories) or ThermoFisher Scientific™ Shandon™ Immuno-Mount™ (Cat. #9990402, ThermoFisher Scientific) mounting medium. Sections from P21 animals were counterstained with DAPI (Cat. # MP-01306, Invitrogen) prior to coverslipping. For *in situ* hybridization, e14.5 and e18.5 were processed as in ^15^, using the *Lhx6* digoxigenin-labeled probe ^113^.

After their respective culture time, electroporated MGE explants were fixed ON at 4°C with 4% PFA, while *Trio^-/-^* and WT MGE explants were fixed for 1h. After several rinses in PBS, explants were blocked for 1h at RT as above, followed by an ON incubation at 4°C in the primary antibody solution (5% normal goat serum, 0.1% Triton X-100) containing 1:1000 dilutions of Rabbit anti-GFP (Cat. #A-6455, ThermoFisher Scientific) or Rat anti-mCherry (Cat. # M11217, ThermoFisher Scientific) antibodies, or a 1:100 dilution of mouse anti-βIII Tubulin (Cat # G7121, Promega) antibody. For F-actin labeling, explants were stained with Alexa Fluor 488-Phalloidin (1:200, Molecular Probes, ThermoFisher Scientific). Explants were then rinsed in PBS several times followed by a 2h incubation at RT in the secondary antibody solution (5% normal goat serum, 0.1% Triton X-100) containing 1:1000 dilutions of the goat anti-Rabbit Alexa 488 and goat anti-Rat Alexa 594 antibodies or a 1:400 dilution of the donkey anti-mouse Alexa 488 or of the donkey anti-mouse Alexa 594 (Jackson ImmunoResearch), together with Hoechst staining (30 min, RT). Then, explants were washed again three times in PBS and were immediately imaged by confocal microscopy.

### EdU staining

Pregnant dams were injected intraperitoneally at e12.5 with a solution containing 5-Ethynyl-2**’**-deoxyuridine (EdU, ThermoFisher) and embryos were collected 2h after. Brains were obtained and fixed as above, cut into 18 μm-thick cryosections and processed following manufacturer instructions (Click-iT EdU Alexa Fluor 488 Imaging kit, Life Technologies) for 30 min at RT. Sections were rinsed three times in 3% bovine serum albumin (BSA) and then in PBS. Hoechst staining was performed for 30 min at RT before pursuing the immunohistochemistry protocol as described above.

### Plasmids

The pIRES2-EGFP vector (Cat. #V11106; Clontech) was modified by enzymatic restriction to insert the Dlx5/6 promoter (Gift from G. Fishell) ^114^ in place of the original CMV promoter. The *Trio* full-length cDNA (*Trio^F-L^*) was cloned by standard PCR using a cDNA library from 3 adult mouse cortices and inserted into the modified Dlx5/6::pIRES2-EGFP vector by enzymatic restriction. The *Trio*^Δ*GEFD*^^1^ and *Trio*^Δ*GEFD*^^2^ plasmids were obtained by mutagenesis overlapping PCR using the *Trio^F-L^* cDNA plasmid as template. Inserted mutations allowed for the deletion of the GEFD1-PH1 domain (Δ1291-1591) or the GEFD2-PH2 domain (Δ1969-2271). The resulting plasmid was inserted into the modified *Dlx5/6::pIRES2-EGFP* vector by enzymatic restriction. The *Dlx5/6::mCherry* plasmid, generated by replacing the EGFP cassette by an *mCherry* cassette, was used as control. The *CMV::tdTomato-Lifeact-7* plasmid (abbreviated Lifeact throughout the manuscript) was a gift from Michael Davidson (Addgene plasmid # 54528; http://n2t.net/addgene:54528; RRID:Addgene_54528) ^115^. All plasmids were verified by sequencing before use.

### Ex utero electroporation

Electroporation of the MGE was conducted at e13.5 as previously described ^112^, with minor modifications. Briefly, endo-free plasmid DNA solutions, mixed with 0.01% Fast Green (Cat. #F7258, Sigma-Aldrich), were injected *ex vivo* directly into the MGE of e13.5 embryos. The *Dlx5/6::mCherry* control plasmid was either mixed with vehicle (TE buffer) or with one of the *Trio* cDNA plasmid to obtain a final concentration of 1.5 µ g/µ l each. The Lifeact plasmid was electroporated at 1.5 µ g/µ l. Electroporation was performed with platinum plated electrodes (Tweezertrodes, BTX Harvard Apparatus) and an Electro Square Porator (BTX Harvard Apparatus) by delivering 4 square pulses of 40 V, 50 ms in duration at 500 ms intervals.

### Organotypic slice cultures and MGE explant cultures

*Trio^WT^* and *Trio^cKO^* embryonic brains were processed for organotypic slice cultures as previously described ^112^. Briefly, e13.5 brains were dissected out of the skull in cold L-15 medium (Invitrogen), embedded in 4% low-melting point agarose and cut with a vibratome into 250 µ m-thick slices collected in sterile oxygenated artificial cerebrospinal fluid (ACSF; see ^112^ for composition). Slices that contained the MGE were cultured on 30 mm membrane inserts in enriched Neurobasal medium (see ^112^ for composition) for 2h before being transferred into an 8-well coverslip for time-lapse imaging. For *Trio^cKO^ and Trio^WT^* MGE explant cultures, cortices from WT (without the Cre and the RCE alleles) e13.5 littermate embryos were dissected out, mechanically dissociated in enriched Neurobasal culture medium ^112^ and plated at a density of 5.25 x 10^5^ cells/cm^2^ ^116^ in 8-well coverslips previously coated with collagen and poly-L-lysine, followed by culture for 1h (37°C, 5% CO_2_). Then, GFP+ littermate embryos were individually processed by decapitation, followed by plasmid injection and MGE electroporation, when applicable, dissection of the MGE and cutting into approximately 200-µ m^2^ explants, which were cultured onto the previously prepared monolayer of mixed cortical cells for 48h. For *Trio^-/-^* and WT MGE explants, e14.5 brains were processed as above to obtain 250-µ m thick coronal sections. Then, MGE were dissected out of the obtained slices, cut into small explants as above using a tungsten needle and either cultured on poly-L-ornithine/laminin coated wells (both from Sigma-Aldrich) for growth cone morphology analysis or embedded in rat tail collagen (BD Bioscience) for migration front analysis. The explants were cultured for 48h in Neurobasal medium supplemented as above, but with the addition of 100 U/ml pen-strep (Invitrogen).

### Time-lapse imaging and video analyses

Migrating INs were monitored by time-lapse imaging of neocortical EGFP+ cells in organotypic slices, EGFP+ cells in MGE explants or mCherry+/EGFP+ cells in electroporated MGE explants. For the migration and branching assays, images were acquired every 5 min for 5h using a Leica spinning disk confocal microscope equipped with an environmental chamber (37°C, 5% CO_2_), a sCMOS Orca Flash 4.0 camera, a HC PL APO 20x/0.80 objective and the VisiView acquisition software (v. 4.1.0.6). Images from e13.5 organotypic slices were taken at the migration front in the dorsal pallium with 18 optical z-plans at 2 µ m intervals. Images of EGFP+ cells migrating out of MGE explants were acquired using the same parameters. Migration dynamics was analyzed according to the method described in ^36^. Briefly, a nuclear translocation was identified when the movement of the nucleus exceeded 5 µ m. For organotypic slices, only tangentially migrating INs were included in the migration analyses. The amplitude of translocation was calculated as the average length of significant nuclear movement per cell. The frequency of translocation corresponds to the number of significant nuclear movement divided by the time of recording (5h). The total distance corresponds to the length of the entire track covered by the soma, whereas the displacement refers to the net distance between the first position (at 0h) and the last position of the soma (at 5h). The global velocity was calculated by dividing the total distance (nm) by the time of recording (s). INs were considered in pause when cell body movement were less than 5 µ m in amplitude, and pauses were calculated in terms of duration and frequency.

Nucleokinesis velocity corresponds to the mean speed of nuclear translocations, excluding pauses. Branching dynamics was analyzed for the leading process only. The growth cone splitting rate was quantified by dividing the number of new branches or growth cone splittings by the time of recording. The transient branch lifetime corresponds to the average time between the appearance of a new branch or a growth cone splitting and either its complete retraction or its transformation into a trailing process. A multipolar phenotype (number of primary neurites > 1) was identified whenever two or more primary branches were simultaneously extending with at least one growth cone each. Secondary neurites were identified as newly formed branch or growth cone splitting closest to the cell body. Given the dynamic nature of migrating INs and the quick switch from leading to trailing process and vice-versa, we considered any new branch or growth cone splitting that formed more distally than a secondary neurite as a tertiary neurite. Quantifications correspond to the total number of such events (primary, secondary, tertiary neurites) for the entire time of recording per cell. For actin dynamics assays, images were acquired every 30 s for 3h using the same system as above, but with a N PL L 40x/0.55 CORR objective, and analyzed using fire-mode in the ImageJ/FIJI software to visualize actin intensity and localization, as in ^36^. Briefly, for each electroporated cell, each timeframe was analyzed and F-actin was categorized as being either diffusely distributed, forming a rear cup behind the nucleus or condensing into a rear dot. The number of event for each category was then divided by the total number of timeframes analyzed to obtain the proportion of time spent in each state.

### Microscopy, neuronal quantifications and morphological analyses

Immunostained coronal 18 µ m-thick sections from e13.5, e15.5 were scanned using an upright Leica TSC SP8 confocal microscope equipped with three solid state lasers, PMT and HyD detectors and the Leica Application Suite X (LAS-X) acquisition software (v. 3.5.19976.5, Leica Microsystems). For each brain section, several z-stacks containing six or seven 2 µ m-thick optical sections were acquired using a HC PL Apo 20x/0.75 dry objective and tiled to cover the entire section. Isolated GABAergic INs at the migratory front in 50 µ m-thick immunostained sections were imaged in their entirety using the same confocal microscope with a 63x/1.40 oil objective, a digital magnification of 1.5 and 0.5 µ m-thick optical sections. Immunostained coronal 18 µ m-thick sections from P21 animals and electroporated INs in MGE explants were imaged with an inverted Leica SP8 confocal microscope, equipped as above, but with four solid state lasers and using a HC PL Apo 20x/0.75 dry objective and a HC PL APO 63x/1.40 oil CS2 objective, respectively. Electroporated INs in MGE explants were imaged in their entirety.

Hybridized e14.5 and e18.5 coronal sections were imaged with a Leica MZ16F Stereoscope. For EdU experiments at e12.5, images were acquired using an Olympus Widefield BX63 microscope equipped with a Hamamatsu ORCA-flash4 camera and a 10x objective. Explants and growth cones from WT and *Trio^-/-^* embryos were imaged with a Leica DMi8 microscope using a HC PL Fluotar 10x/0.30 objective also equipped with a Hamamatsy ORCA-flash4 camera. For illustration purposes only, growth cones were also imaged using a HC PL Fluotar 20x objective.

Neuronal quantifications in embryonic brains were performed using an unbiased approach with the open source Fiji/ImageJ software. Eight equally spaced sections were imaged in tile-scans and z-stacks across the entire rostro-caudal extent of the embryonic telencephalon, with the first section selected randomly. For each coronal section, three optical sections in the middle of the stack were selected and processed by average intensity projection. On the resulting images, the dorsal pallium was then divided in five equal bins across its entire ventro-dorsal extent, starting at the cross-section between the dorsal pallium and the LGE and GFP+ cells were quantified in each bin, as in ^117^. For e15.5 and e18.5 brains, the dorsal pallium was divided in ten equally distant bins in the latero-medial plan, and embryonic rudimental layers, i.e. the marginal zone (MZ), the cortical plate (CP), the intermediate (IZ), subventricular (SVZ) and ventricular zones (VZ) were defined, as in ^117^. GFP+ cells were manually counted in each of these bins and/or layers and their percentile distribution across all bins was reported. For the quantification of radially migrating cells, the number of GFP+ cells oriented 45° or more from the SVZ and MZ migration streams was quantified and reported as a proportion of the total number of GFP+ cells in the dorsal pallium (e13.5) or in the CP (e15.5). For Lhx6 signal quantification, signal intensity in each bin was measured and normalized to the total signal intensity detected in all bins. The resulting signal intensity percentage was considered proportional to the number of cells stained in each bin. For proliferation assays at e12.5, the number of EdU-positive cells in the VZ, SVZ and mantle zone was calculated and normalized to the total number of Hoechst-positive cells. For P21 brains, GFP-positive INs were manually counted in the somatosensory cortex using Fiji/ImageJ software as in ^61^. The cortical layers were identified using DAPI staining. Briefly, fate-mapped GABAergic INs were identified as such by their expression of EGFP and DAPI, and were manually counted using the Cell Counter plug-in in Fiji/ImageJ. Densities were calculated by dividing the number of cells by the area of each layer, or by the total area. A similar approach was used to quantified immunostained INs expressing either PV, SST, CR or VIP, with the exception that only immunostained cells that also co-expressed EGFP were considered for quantification.

Morphological analyses were performed on high-resolution confocal images using the specialized NeuroLucida image analysis software (v. 11.0.8.2, MicroBrightField) to reconstitute neurons in 3-D. The cell body size corresponds to the largest area of the cell body. The leading process was identified by the presence of a swelling in front of the cell body, the thickness of the neurite and the presence of growth cones at the extremities. The trailing process was identified as such by the presence of a thin neurite, generally oriented towards the back of the cell body, and devoid of a growth cone or a swelling. Migration distances in fixed MGE explants generated at e14.5 were obtained by measuring the mean distance of the 50 cells that migrated the most from the explant and averaged between at least four different explant per brain. Growth cones were scored as in ^118^. Briefly, growth cones were categorized as either collapsed or with a complex morphology, and for growth cones presenting with a complex morphology, the surface was measured using the area tool in Fiji/ImageJ.

### Statistical analyses

All statistical analyses were performed using GraphPad Prism software (v. 8.2.0, GraphPad software, San Diego, USA). Unpaired Student’s *t*-tests were used for comparisons between two groups with normal distribution, while Mann-Whitney tests were used for comparison between two groups with non-parametric data distribution or with small samples. One-way ANOVA, followed by Tukey’s multiple comparisons tests, were used for comparisons between three or more groups and two-way ANOVA or two-way mixed-effect analyses (for repeated measures with missing values), followed by Bonferroni’s, Sidak’s or Tukey’s multiple comparisons tests were used to compare groups with two or more factors. Fisher’s exact test was used for growth cone morphology comparison. Differences were considered significant at *P* < 0.05. Mean and standard error of the mean (s.e.m.) are used throughout the text as central tendency and dispersion measures, respectively.

Supplementary information is available at MP’s website.

## Figure Legends

**Supplementary Figure 1. Defective laminar distribution of *Lhx6+* cortical interneurons in *Trio^-/-^* embryos**. (**A-D**) *Lhx6 in situ* hybridization on e14.5 (**A-B**) and e18.5 (C-D) coronal brain sections in WT (**A, C**) and *Trio^-/-^* (**B, D**) embryos. (**E**) Histogram showing the distribution of *Lhx6+* signal in the dorsal pallium of e14.5 WT and *Trio^-/-^* embryos. Bins are delineated by the dotted lines in **A-B** (n=5 WT and 3 *Trio^-/-^* embryonic brains). (**F**) *Lhx6+* signal distribution within the e18.5 somatosensory neocortex layers in wild type (**C**) and *Trio^-/-^* (**D**) (n=3 WT and 3 *Trio^-/-^* embryonic brains). (**G-K**) Reduced migration of MGE-derived cells in *Trio-/-* cultured MGE explants. (**G-J**) Representative immunofluorescence images of WT (**G, H**) and *Trio^-/-^* (**I, J**) MGE explants generated at e14.5 and cultures for 48h. (**K**) Histogram showing the mean maximal distance traveled by WT and *Trio^-/-^* MGE-derived cells in cultured explants (n= 11 explants from 2 brains for each genotype). **(L-Q)** Impaired growth cone morphology in *Trio^-/-^* MGE-derived cells. (**L-O**) Representative illustrations for growth cone complex morphology (**L, N**) and collapsed morphology (**M, O**) in WT (**L, M**) and *Trio^-/-^* (**N, O**) MGE-derived cells from cultured MGE explants. (**P**) Histogram showing the proportion of WT and *Trio^-/-^* MGE-derived cells with a complex or a collapsed morphology (n = 766 growth cones from 3 WT brains and 565 growth cones from 3 *Trio^-/-^* brains). (**Q**) Histogram showing the mean growth cone surface of WT and *Trio^-/-^* MGE-derived cells with a complex morphology (n = 181 growth cones from 3 WT brains and 65 growth cones from 3 *Trio^-/-^* brains). \**P* < 0.05; \*\**P* < 0.001 \*\*\*\**P* < 0.0001 by two-way ANOVA followed by Sidak’s multiple comparison’s test (**E-F**), or by Mann-Whitney test **K, Q** and Fisher’s exact test for **P** (using the number of growth cones in each category). Data are represented as mean ± SEM. CP, Cortical Plate; IZ, Intermediate Zone; UL, Upper Layers. Scale bars: 500 μm (**A, B**), 250 μm (**C, D**) 150 μm (**G-J**) and 10 μm (**L-O**).

**Supplementary Figure 2: Proliferation in the MGE is not affected in *Trio^-/-^* embryos.** (**A-B**) EdU incorporation in proliferative cells in the MGE at e12.5 in WT (**A**) and *Trio^-/-^* (**B**) embryos. (**C**) Histogram showing the quantification of EdU incorporation in VZ, SVZ and MZ in *Trio^-/-^* mutants compared to wild type littermates (n = 3 WT and 3 *Trio^-/-^* brains). Comparisons were made by two-way ANOVA followed by Sidak’s multiple comparison’s test. Data are represented as mean ± SEM. MZ, mantle zone; SVZ, subventricular zone; VZ, ventricular zone. Scale bar: 250 μm.

**Supplementary Figure 3. *Trio^cKO^* INs in MGE explant cultures display migration deficits as in organotypic brain slice cultures.** (**A**) Scheme of the experimental procedure for live-imaging migrating INs in MGE explant cultures. (**B-C**) Detailed time-lapse sequences of GFP+ MGE INs (arrowheads) from *Trio^WT^* (**B**) and *Trio^cKO^* (**C**) brains migrating in MGE explants cultures over the course of 5h. (**D**) Histograms of the migration dynamics of *Trio^WT^* and *Trio^cKO^* GFP+ INs in MGE explant cultures, including the velocity, total distance, net displacement and persistence ratio (directionality). (**E**) Histograms showing more detailed nucleokinesis dynamics of *Trio^WT^* and *Trio^cKO^* GFP+ INs in MGE explant cultures, including the velocity, frequency, duration and amplitude of nuclear translocation. (**F**) Histograms showing the quantification of migrations pauses during the migration of *Trio^WT^* and *Trio^cKO^* GFP+ INs in MGE explant cultures, including the frequency and duration of pauses (n = 81 cells from 3 *Trio^WT^* brains and 56 cells from 3 *Trio^cKO^* brains). (**G**) Scheme for live-imaging of electroporated *Trio^WT^* and *Trio^cKO^* GFP+ INs in MGE explant cultures. (**H-K**) Histograms showing the total distance (**H**), nucleokinesis frequency (**I**), migration pause frequency (**J**) and migration pause duration (**K**) following electroporation of the control mCherry plasmid in *Trio^WT^* or *Trio^cKO^* INs or a modified *Trio* cDNA in *Trio^cKO^* INs, as in Figure 5. (**L-O**) Histograms of branching dynamics showing the mean branch lifetime (**L**), as well as the number of (**M**) primary, (**N**) secondary and (**O**) tertiary neurites following electroporation of the above-mentionned plasmids. \**P* < 0.05, \*\*\**P* < 0.001 and \*\*\*\**P* < 0.0001, by Student’s t-test (**D-F**) or one-way ANOVA followed by Tukey’s multiple comparisons test (**H-O**). Note that for better clarity, only comparisons with *Trio^cKO^* INs electroporated with mCherry (black column) are shown in **H-O**, but *p*-values for all comparisons can be found in Supplementary Table 1. Scale bar: 50 µ m.

**Supplementary video 1.** Example of a time-lapse recording for an e13.5 *Trio^WT^* organotypic brain slice after 2h of culture.

**Supplementary video 2.** Example of a time-lapse recording for an e13.5 *Trio^cKO^* organotypic brain slice after 2h of culture.

**Supplementary video 3.** Example of a time-lapse recording for *Trio^WT^* MGE-INs migrating out of an MGE explant generated at e13.5 and cultured for 48h.

**Supplementary video 4.** Example of a time-lapse recording for *Trio^cKO^* MGE-INs migrating out of an MGE explant generated at e13.5 and cultured for 48h.

**Supplementary video 5.** Example of a time-lapse recording for *Trio^WT^* MGE-INs (green) electroporated with the *Dlx5/6::mCherry* control plasmid (red) and migrating out of an MGE explant generated at e13.5 and cultured for 48h. Electroporated MGE-INs appear in yellow.

**Supplementary video 6.** Example of a time-lapse recording for *Trio^cKO^* MGE-INs (green) electroporated with the *Dlx5/6::mCherry* control plasmid (red) and migrating out of an MGE explant generated at e13.5 and cultured for 48h. Electroporated MGE-INs appear in yellow.

**Supplementary video 7.** Example of a time-lapse recording for *Trio^cKO^* MGE-INs (green) electroporated with *Trio*^Δ^*^GEFD1^* (red) and migrating out of an MGE explant generated at e13.5 and cultured for 48h. Electroporated MGE-INs appear in yellow.

**Supplementary video 8.** Example of a time-lapse recording for *Trio^WT^* MGE-Ins electroporated with the Lifeact plasmid and migrating out of an MGE explant generated at e13.5 and cultured for 48h. For illustration purposes, the electroporated MGE-IN appears in fire mode.

**Supplementary video 9.** Example of a time-lapse recording for *Trio^cKO^* MGE-INs electroporated with the Lifeact plasmid and migrating out of an MGE explant generated at e13.5 and cultured for 48h. For illustration purposes, the electroporated MGE-IN appears in fire mode.

**Supplementary Table 1.** Summary data table comprising the mean values, standard errors to the mean as well as p-values for each experiment and each genotype.

## Notes

### Competing Interest Statement

The authors have declared no competing interest.

