## Supplementary figures and images for "Both GEF domains of the autism and epilepsy-associated Trio protein are required for proper tangential migration of GABAergic interneurons"

### Supplemental figure 1

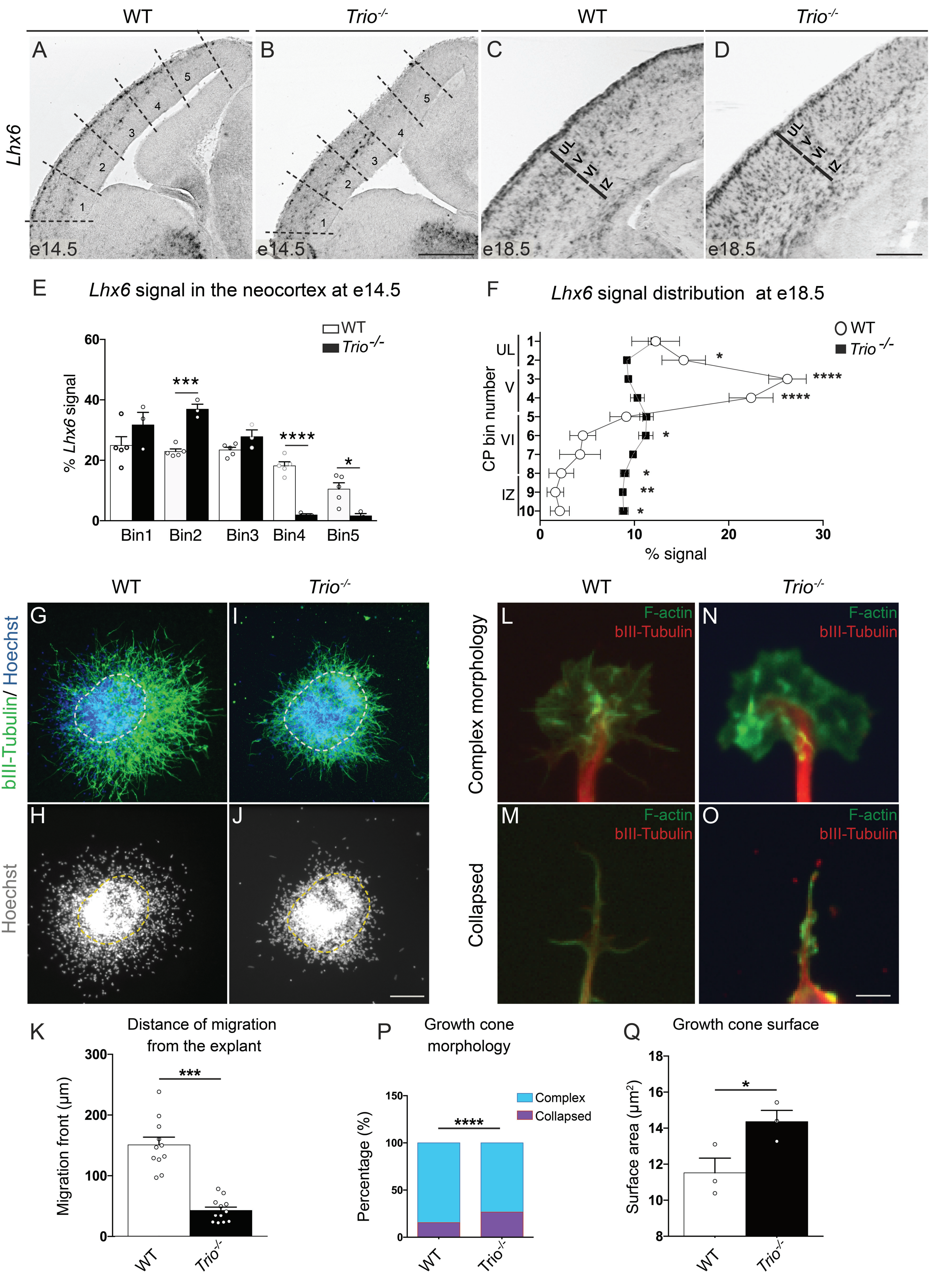

### Supplemental figure 2

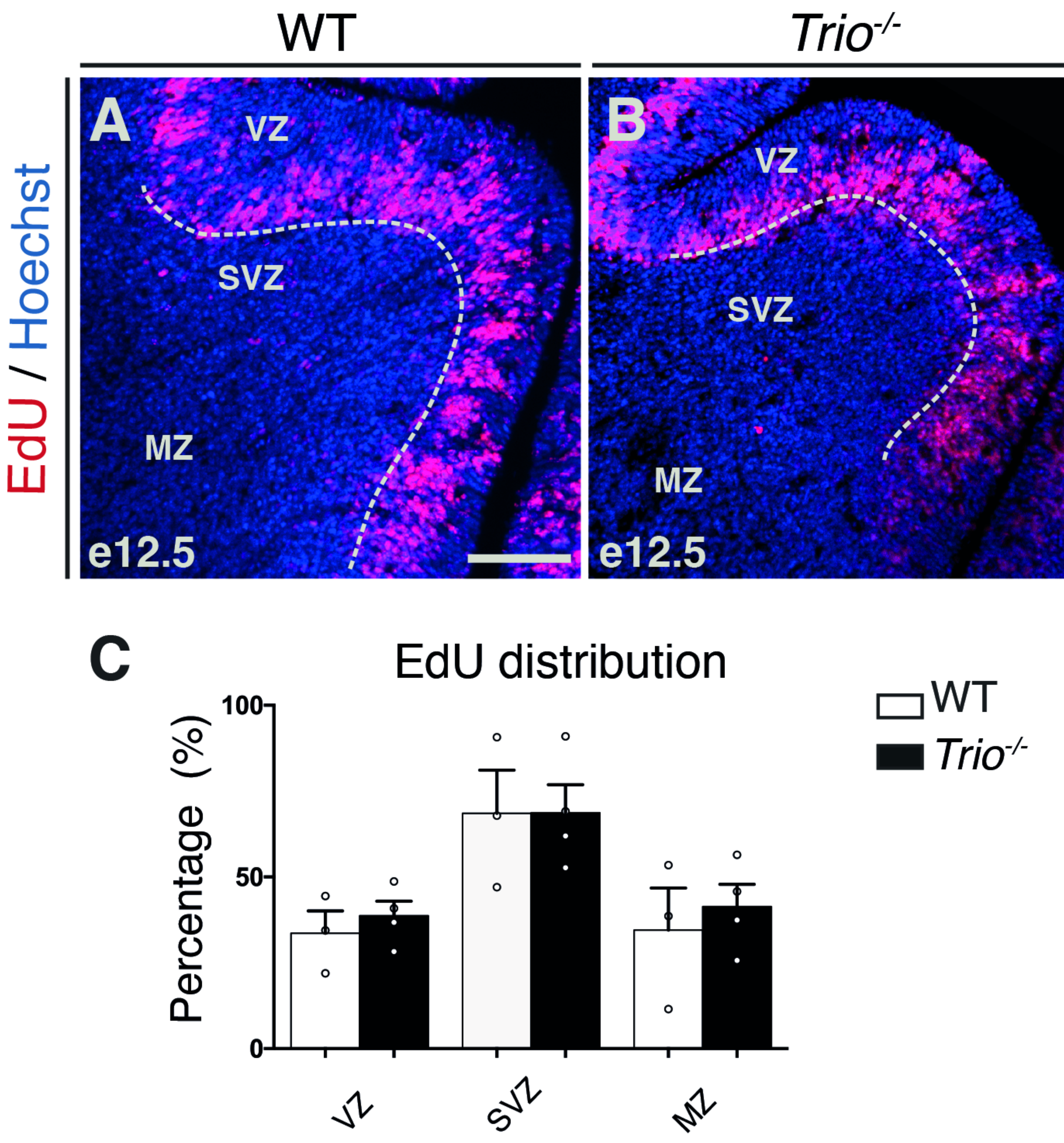

### Supplemental figure 3

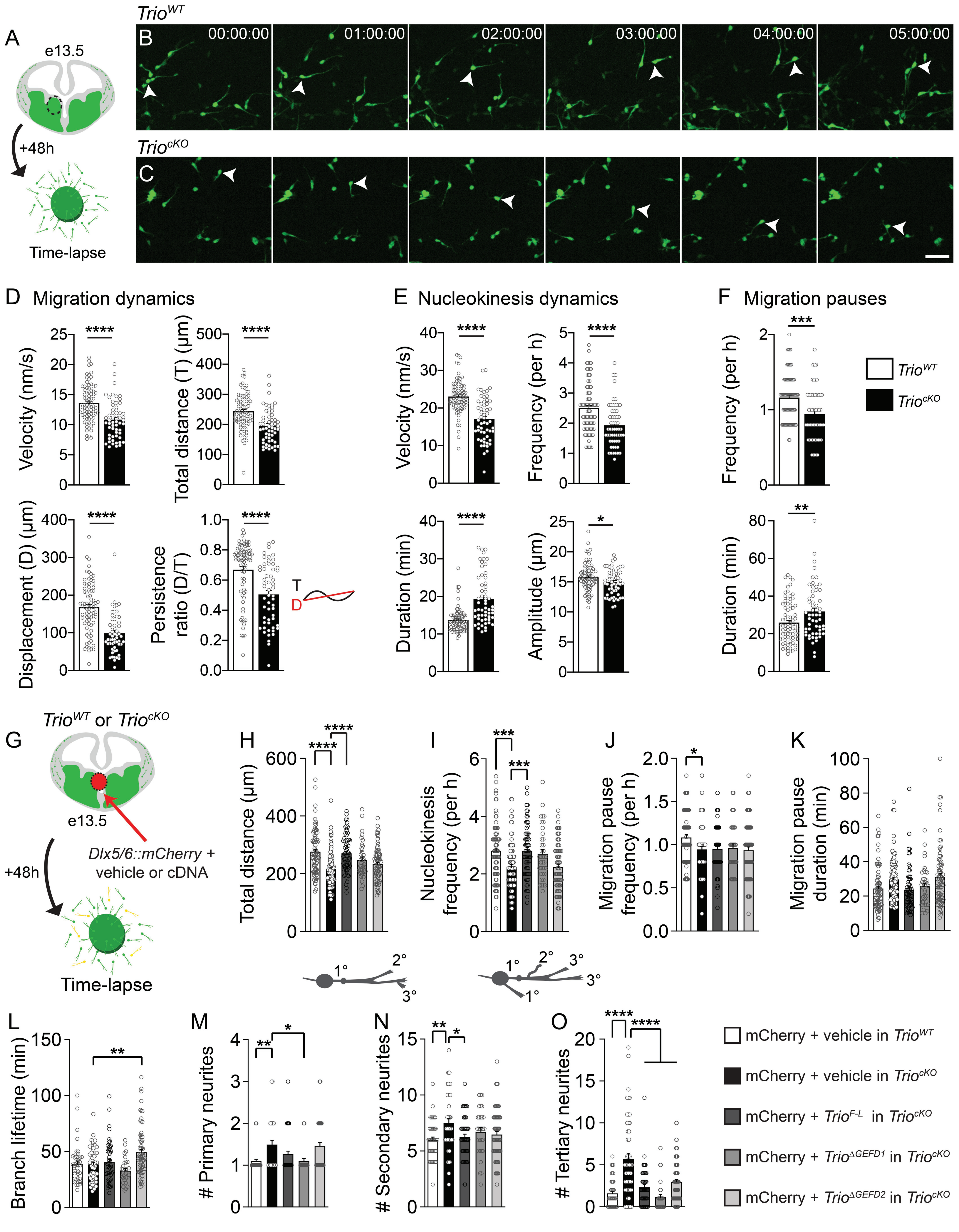
